# Spatial immune hubs defined by conserved activated dendritic cells are remodeled by immunotherapy

**DOI:** 10.64898/2026.02.10.703136

**Authors:** Tomoyuki Minowa, Sunita Keshari, Shan He, Akata Saha, Alexander S. Shavkunov, Josué E. Pineda, Yao Yu Yeo, Bokai Zhu, Vakul Mohanty, Sonali Jindal, Ken Chen, Kenneth H. Hu, Linghua Wang, Garry P. Nolan, Stephanie S. Watowich, Matthew M. Gubin

**Affiliations:** Department of Immunology, The University of Texas MD Anderson Cancer Center, Houston, TX, USA; Department of Bioinformatics and Computational Biology, The University of Texas MD Anderson Cancer Center, Houston, TX, USA; Department of Medical Oncology, Dana-Farber Cancer Institute, Boston, MA, USA; Ragon Institute of MGH, MIT, and Harvard, Cambridge, MA, USA; Broad Institute of MIT and Harvard, Cambridge, MA, USA; Massachusetts Institute of Technology, Cambridge, MA, USA; James P. Allison Institute, The University of Texas MD Anderson Cancer Center, Houston, TX, USA; Department of Genomic Medicine, The University of Texas MD Anderson Cancer Center, Houston, USA; The University of Texas MD Anderson Cancer Center UTHealth Houston Graduate School of Biomedical Sciences, Houston, USA; Institute for Data Science in Oncology, UT MD Anderson Cancer Center, Houston, TX, USA; Department of Pathology, Stanford University, Stanford, CA, USA; Department of Microbiology and Immunology, Stanford University, Stanford, CA, USA; Platform for Innovative Microbiome and Translational Research, MD Anderson Cancer Center, Houston, TX, USA

**Author notes:** **Correspondence**: Matthew M. Gubin, PhD and Stephanie S. Watowich, PhD, Department of Immunology, The University of Texas MD Anderson Cancer Center, 7455 Fannin St., Houston, TX 77054. These authors contributed equally (Tomoyuki Minowa, Sunita Keshari). Present address: Department of Experimental Radiation Oncology, The University of Texas MD Anderson Cancer Center, Houston, TX, USA. Present address: Computational Sciences Center of Excellence, Department of Computational Biology and Medicine, Genentech Research and Early Development, Genentech, CA, USA.

## Abstract

Dendritic cells (DCs) orchestrate anti-tumor immune responses, yet the full extent of their phenotypic diversity, and spatial dynamics within the tumor microenvironment (TME) remains incompletely understood. Here, we constructed an integrated atlas of tumor-infiltrating DCs by harmonizing single-cell transcriptomic data from 12 murine tumor studies and 28 published human cancer datasets, together with newly generated single-cell-resolved multiplexed tissue imaging across immunotherapy conditions in a murine model. We noted conserved transcriptional states across species, including canonical conventional type 1 DCs (cDC1s), diverse type 2 DC (cDC2) subpopulations, and two activation states characterized by CCR7 expression (CCR7^+^ DCs) or interferon-stimulated gene expression (ISG⁺ DCs). Spatial transcriptomics analyses from human TMEs revealed that CCR7^+^ DCs and ISG⁺ DCs reside in distinct T cell-enriched regions that are embedded within distinct signaling environments. High-dimensional multiplexed proteomic imaging demonstrated that these DC-T cell niches undergo divergent remodeling across multiple immunotherapy conditions. Notably, this spatial reorganization occurred despite minimal detectable changes in DC transcriptional states. This study delineates conserved DC activation states and their spatial organization within tumors and captures the therapy-dependent remodeling, providing a framework for studying therapy-associated remodeling of DC immune programs in cancer.

## INTRODUCTION

Dendritic cells (DCs) are pivotal mediators of both innate and adaptive immune responses(*1–5*). DCs comprise multiple subtypes characterized by distinct ontogeny, transcriptional programs, and functional properties, including plasmacytoid DCs (pDCs), conventional DCs (cDCs), and monocyte-derived DCs (moDCs)(*6*). As the name implies, moDCs arise from circulating monocytes and are predominantly associated with inflammatory settings, whereas pDCs are specialized for their abundant production of type I interferons (IFN-Is) in response to viral infections(*7*).

cDCs are broadly subdivided into two main subsets: type 1 cDCs (cDC1s) and type 2 cDCs (cDC2s), each cDC regulated by distinct transcription factors and characterized by discrete phenotypic and functional features(*1–5*). As professional antigen-presenting cells (APCs), cDCs are capable of processing and presenting both exogenous and endogenous antigens on MHC class I and MHC class II molecules, respectively, thereby enabling the activation of CD8 and CD4 T cells. This function is central to anti-tumor immunity, as cDCs capture tumor-derived antigens and initiate T cell priming within tumor-draining lymph nodes (tdLNs) together with co-stimulatory molecules. In addition, tumor-infiltrating cDCs further regulate these primed T cells locally within the tumor microenvironment (TME). Beyond antigen presentation, soluble factors produced by tumor-infiltrating cDCs contribute to T cell activation, function, and survival (e.g., IL-12 and IL-15) and promote the recruitment of T cells into DC-rich niches through chemokine production (e.g., CXCL9 and CXCL16)(*2, 4*). These DC–T cell interactions occur within spatially organized niches, where additional immune cell populations further shape T cell responses. For example, CD4 T cells have been shown to support CD8 T cell responses through multiple mechanisms. Through these coordinated processes, cDCs shape T cell activation, proliferation, trafficking, and effector differentiation and functions. Importantly, the presence and organization of such DC–T cell niches have been implicated in effective anti-tumor immune responses and are increasingly recognized as key determinants of responsiveness to cancer immunotherapy(*8–11*).

Although cDC1s and cDC2s are well defined at homeostasis based on marker expression and transcriptional signatures, the tumor context introduces additional layers of heterogeneity that give rise to distinct DC activation states(*1*). For instance, one such state is a population of migratory CCR7⁺ DCs enriched within the TME. These cells are marked by expression of FSCN1 and LAMP3, maturation-associated markers such as CD40 and CD83, and multiple immunoregulatory molecules, including PD-L1, PD-L2, and CD200(*12–16*). These DCs have been described using diverse nomenclature across studies, including “DC3s”(*15, 17, 18*), “mature DCs enriched in immunoregulatory molecules” (mregDCs)(*12*), CCR7^+^ DCs(*19*) or LAMP3⁺ DCs(*14*). Hereafter, we will refer to this population as CCR7^+^ DCs.

Another activation state comprises DCs marked by a transcriptomic profile enriched for IFN-stimulated genes (ISGs), termed ISG⁺ DCs. This phenotype has been described within both cDC1s(*9, 20, 21*) and cDC2s(*14, 22, 23*) compartments. In preclinical mouse models, ISG⁺ DCs were shown to promote robust T cell-mediated antitumor immunity and are required for optimal responses to immune checkpoint blockade(*23*). However, the conservation of these activation states across cancer types and species, as well as their spatial localization within the TME, and their role in immune checkpoint therapy (ICT) response, remain poorly defined.

In this study, we constructed an atlas of tumor-infiltrating DCs by integrating multimodal data comprising large-scale single-cell RNA sequencing (scRNA-seq) datasets from human and murine tumors and spatial transcriptomics from human colorectal cancer with newly generated, single-cell-resolved multiplexed tissue imaging that includes distinct ICT conditions in a murine melanoma model. By focusing on spatial organizations and therapy-associated alterations of DCs emphasizing two DC activation states (CCR7^+^ DCs and ISG⁺ DCs) within the TME, we provide novel insights into conserved DC activation states and their potential biological and clinical relevance in cancer immunotherapy.

## RESULTS

### Construction of Tumor-infiltrating DC Signatures Across Human Cancer Types

To characterize the cellular phenotype of tumor-infiltrating DCs, we first compiled 28 scRNA-seq datasets derived from human solid tumors spanning 12 distinct cancer types and normal tissues that included DC populations (**Supplemental Figures 1A, B, Supplemental Table S1, and see Methods**). For this study, we chose to focus on cDCs and thus, we excluded pDCs from subsequent analyses. This analysis yielded a single-cell DC atlas (exclusive of pDCs) consisting of 29,554 cells including 17,092 tumor-infiltrating DCs from 353 samples representing 12 tissues (**Figure 1A, and Supplemental Figures 1C–F**). Unsupervised clustering of DCs identified eight clusters, which were further classified into three major DC types based on differentially expressed genes (DEGs) and established canonical markers(*24*): cDC1 (*XCR1, CADM1, CLEC9A, BATF3*), cDC2/cDC2-like (*CD1C, CLEC10A, SIRPA*), and CCR7^+^ DCs (*LAMP3, CCR7, FSCN1, CD40, CD274*), also commonly referred to as mregDCs or LAMP3^+^ DCs (**Figures 1A–C, Supplemental Figure 1G, H, and Supplemental Table S2**). Among six cDC2/DC2-like clusters, cells in C0, C1, C6, and C7 exhibited high expression of *IL1B*, which encodes the pro-inflammatory cytokine IL-1β (**Figure 1C**), suggesting that they correspond to the previously reported inflammatory DC2s (inf-DC2s)(*14*). Among these, C1_inf-DC2 and C7_inf-DC2 expressed high amounts of *CCL3* and *CCL4*, chemokines known to be produced by activated DCs that recruit CCR5⁺ CD8 T cells(*25*) (**Figure 1C**). In addition, a proportion of C7_inf-DC2 expressed *CD1A* and *CD207*, markers typically associated with Langerhans cells (**Figure 1C**). Unlike bona fide Langerhans cells in the epidermis, C7_inf-DC2 were enriched in cancer tissues and distributed across various cancer types (**Figures 1D, E**), suggesting that they represent CD207⁺ cDC2s, which have been reported to be present in the TME(*3*). C6_inf-DC2 expressed high levels of monocyte-related markers such as *CD14*, *CD163*, *FCN1*, and *S100A9*, while also expressing *FLT3* and being observed in the TME and cancer-adjacent tissue as well as in blood of healthy donors (**Figures 1B–D**). This cluster resembled circulating inflammatory CD1C⁺CD163⁺ DCs, which have been proposed to arise from monocyte-DC progenitors (MDP) independently of common DC progenitors (CDP) and common monocyte progenitors (cMoP)(*26–29*). C0_inf-DC2 comprised a population exhibiting higher expression of *CCL22* and *CD86* compared with other inf-DC2s and was enriched in the TME and adjacent tissues (**Figures 1C, D**). C3_DC2 was observed in both the spleen and blood and displayed low to intermediate levels of cytokine transcripts and markers associated with DC maturation/activation (*CD80*, *CD83*, *CD86*) (**Figures 1C, D**), suggesting a non-activated state. C5_ISG⁺ DC cluster expresses canonical cDC2 markers, and shows enrichment of IFN-α and IFN-γ signaling pathways, along with increased JAK-STAT pathway activity (**Figure 1C and Supplemental Figures 1I–K**). These findings suggest that C5_ISG⁺ DCs represent an IFN-exposed activation state of cDC2s, a previously described subset(*14, 23*).

**Fig. 1.**
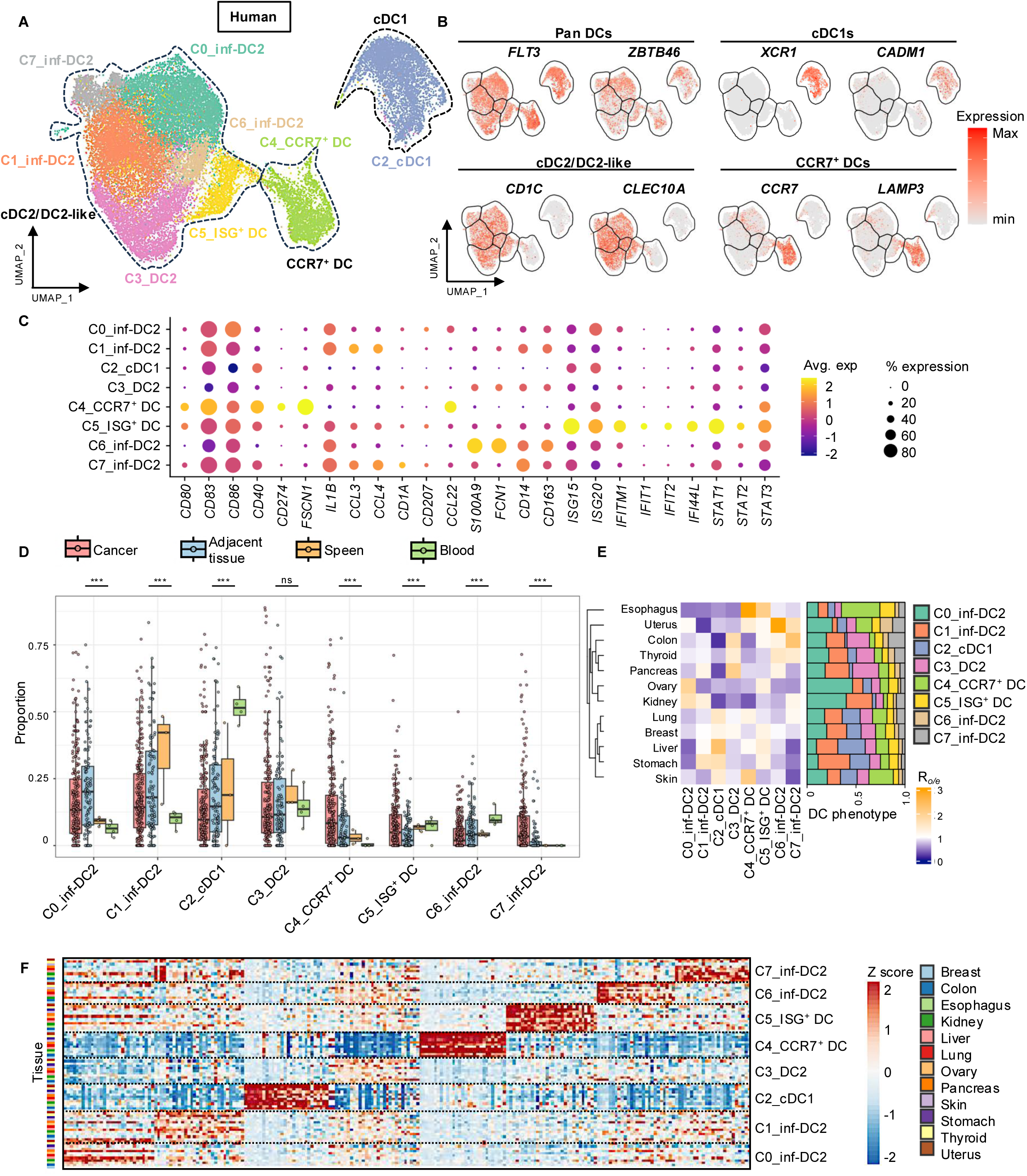
Single-cell transcriptome profiling of tumor-infiltrating DCs in human. **(A)** Integrated UMAP analysis of DCs derived from tumor, adjacent normal tissue, spleen, and blood. Clusters are denoted by colors. **(B)** UMAP analysis of tumor-infiltrating DCs depicting the single-cell expression of selected lineage and phenotypic transcripts. The color scale represents an average expression. **(C)** Dot plot showing the expression of selected signature genes in tumor-infiltrating DCs. Dot size represents the fraction of cells expressing each gene, and color intensity indicates the average expression levels. **(D)** Boxplots showing the fractions of DC subtypes. Spleen and blood samples are from cancer patients and healthy donors, respectively. Each dot represents an individual sample. Samples containing fewer than five cells were excluded from the analysis. The lines within the box and whiskers indicate the median and quartiles, respectively. *P*-values (\**P* < 0.05, \*\**P* < 0.01, \*\*\**P* < 0.001, \*\*\*\**P* < 0.0001) were calculated using one-way ANOVA with Tukey’s multiple comparison test. ns: not significant. **(E)** Heatmap showing tissue preference of each tumor-infiltrating DC cluster, as determined by the ratio of observed to expected cell numbers (R*o/e*). **(F)** Heatmap showing normalized gene expression in tumor-infiltrating DC clusters across cancer types. The heatmap displays the top 30 DEGs with the highest log_2_ fold change and a detection percentage (pct.1) above 0.5, as identified by DEG analysis.

Consistent with prior observations(*14*), cDC2/DC2-like cells were the most abundant DC subtype across cancer types (**Supplemental Figure 1L**). Moreover, cDC2/DC2-like cells exhibited greater heterogeneity than other DC subtypes based on ROGUE score(*30*) (**Supplemental Figure 1M**). Overall, all clusters were broadly distributed across cancer types in the integrated data, and heterogenous cDC2 clusters were consistently preserved across alternative data integration strategies, as evidenced by the retention of distinct cluster identities in UMAP space (**Figure 1E and Supplemental Figure 1N, O**).

### Conserved DC States Across Human Cancer

Previous analyses suggest conservation of cDC1 and CCR7^+^ DC transcriptional programs, with comparatively less conservation observed for the cDC2 phenotype across five human cancer types (lung, breast, liver, colorectal, and ovary)(*16*). To expand upon these findings, we further assessed the extent of conservation using our integrated datasets spanning multiple additional cancer types. C2_cDC1, C5_ISG⁺ DCs, and C4_CCR7^+^ DCs exhibited consistent transcriptional profiles across multiple cancer types; similar conservation was observed for C6_inf-DC2 and C7_inf-DC2, albeit to a lesser extent (**Supplemental Table S3 and Figure 1F**). In contrast, other cDC2/DC2-like clusters displayed less conserved transcriptional patterns (**Figure 1F**). Signature score analysis supported these findings (**Supplemental Figure 1P**). Together, these analyses indicate that cDC1, ISG⁺ DCs, and CCR7^+^ DC transcriptional programs are broadly conserved across human cancers, whereas the cDC2/DC2-like compartment includes both conserved subsets shared across tumor types and subsets with transcriptional features that vary by cancer context.

### Limited Impact of Sex on DC Phenotypes

In recent years, sex-based differences in immune cell phenotype and function have become increasingly recognized. We therefore assessed whether DC phenotypes were potentially influenced by sex. Sex information was available for 63.2% (male: 38.6%, female: 61.4%) of patients and no sex-specific clusters were identified (**Supplemental Figures 2A–D**). After excluding female-overrepresented (breast, uterus, ovary) and male overrepresented (renal) cancers to avoid confounding effects due to highly unbalanced sex representation, DEG analysis revealed that C2_cDC1, C5_ISG⁺ DC, and C4_CCR7^+^ DC exhibited only a small number of DEGs, all of which were restricted to the X or Y chromosomes (**Supplemental Figure 2E**). In contrast, cDC2/DC2-like showed a higher number of DEGs distributed across chromosomes beyond the XY chromosomes. However, cancer-type-specific analyses failed to identify these genes as DEGs, further supporting heterogeneity within cDC2/DC2-like subset across different cancer types (**Supplemental Figure 2F**).

### Conservation of DC Subsets Across Human and Murine Tumors

We next sought to determine whether transcriptional conservation of tumor-infiltrating DC subsets extends across species and tumor contexts. For this, we compiled 12 scRNA-seq datasets from both published and unpublished murine tumor studies, employing the same computational pipeline as the one applied to the human dataset (**Supplemental Figures 3A, B and Supplemental Table S4**). This integrated murine scRNA-seq dataset, comprising 6,592 tumor-infiltrating DCs (excluding pDCs), underwent unsupervised clustering, revealing 11 DC clusters (**Figure 2A and Supplemental Figures 3C–E**). Based on the expression of canonical markers, these clusters were mainly annotated as cDC1 (*Xcr1*, *Cadm1*, *Clec9a*), cDC2/DC2-like (*Sirpa*, *Clec10a*, *Itgam*), and CCR7^+^ DC (*Ccr7*, *Fscn1*, *Cd274*) (**Figure 2B, Supplemental Figure 3F, and Supplemental Table S5**). Similar to the findings in humans, cDC2/DC2-like were more dominant in the TME and exhibited greater heterogeneity compared to other DC subsets (**Supplemental Figures 3G–I**). Analysis of transcriptional profiles across the murine dataset revealed conserved expression patterns in C4/C10_cDC1, C0/7/8/9_ CCR7^+^ DC, and C6_ISG⁺ DC, closely mirroring the observations in the human analysis (**Supplemental Figure 3J**). Notably, direct comparison of human and murine DC clusters demonstrated that the transcriptional profiles of cDC1 and CCR7^+^ DC are well-conserved across species (**Figure 2C**), consistent with prior analyses of human and mouse lung and colorectal cancers(*12, 15, 31*). Further, C6_ISG⁺ DC also displayed conserved patterns across murine and human datasets (**Figure 2C**).

**Fig. 2.**
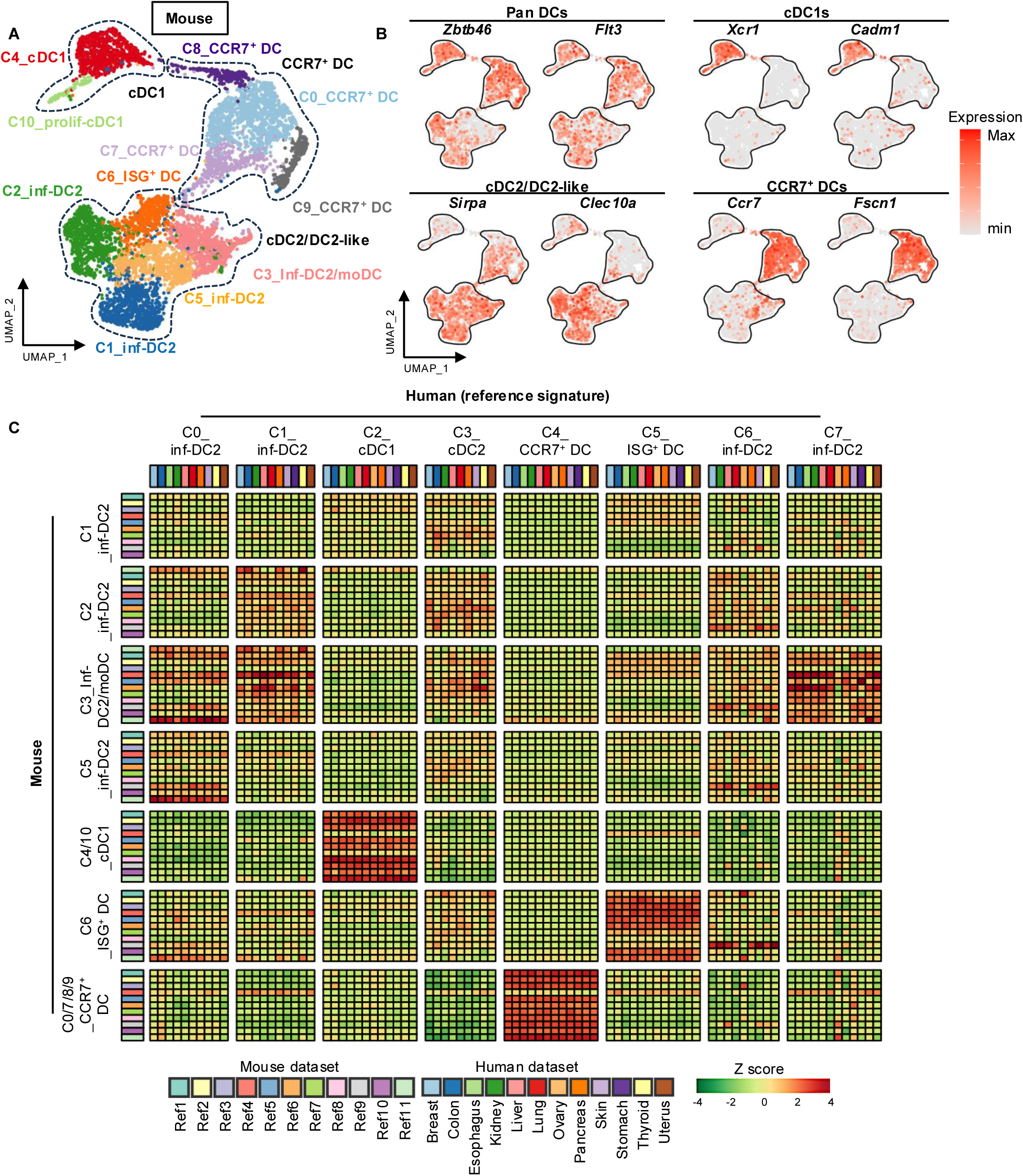
Single-cell transcriptome profiling of tumor-infiltrating DCs in mouse. **(A)** Integrated UMAP analysis of tumor-infiltrating DCs in mouse. Clusters are denoted by colors. **(B)** UMAP analysis of tumor-infiltrating DCs depicting the single-cell expression of selected lineage and phenotypic transcripts. The color scale represents an average expression. **(C)** Heatmap displaying signature scores for tumor-infiltrating DC states across species. Gene signatures for each DC state were determined by DEG analysis of human tumor-infiltrating DCs for each cancer type. These gene signatures were applied to mouse tumor cohorts to calculate cross-species DC signature scores.

### Cross-study Analysis of CCR7^+^ DCs and ISG⁺ DCs

Multiple terminologies have been proposed to describe functionally related DC states, including mregDCs, migratory DCs, and LAMP3⁺ DCs. To contextualize our DC annotations within this framework, we compared our CCR7⁺ DC and ISG⁺ DC clusters with DC subsets defined in previous studies. Cells annotated as LAMP3⁺ DCs by Cheng et al.(*14*) mapped predominantly to our C4_CCR7⁺ DC cluster, whereas our C5_ISG⁺ DC cluster corresponded to cDC2 populations in that study (**Supplemental Figure 4A**). Consistently, our C4_CCR7⁺ DC cluster exhibited high signature scores for CCR7⁺ DCs as defined by Qian et al.(*19*), migratory DCs as defined by Duong et al.(*23*), and mregDCs as defined by Maier et al.(*12*), based on gene signatures derived from these prior reports (**Supplemental Figure 4B and Supplemental Table S6**). These findings indicate that DC subsets annotated across different studies as CCR7⁺ DCs, LAMP3⁺ DCs, or mregDCs share substantial transcriptional similarity, consistent with prior observations(*14, 16*). In contrast, our C5_ISG⁺ DC cluster was enriched for gene signatures previously associated with cDC2-related states, including ISG-DCs and CXCL9⁺ cDC2s, and did not overlap with gene signatures characteristic of moDCs, mregDCs, LAMP3⁺ DCs, or CCR7⁺ DCs (**Supplemental Figure 4B**).

### Transcriptional Links Between ISG⁺ DCs and CCR7⁺ DCs

Although the precise relationship between ISG⁺ DCs and CCR7^+^ DCs remains unclear, the transient enrichment of an ISG transcriptional program was suggested in cDC1s, followed by its subsequent downregulation in mature cDC1s(*20, 21*). Given that the C5_ISG⁺ DC population shares transcriptional features more closely aligned with cDC2s (**Supplemental Figures 5A, B**), we sought to identify the ISG⁺ cDC1 counterpart population. For this, we performed sub-clustering analysis of C4_CCR7^+^ DC and revealed subsets of cells co-expressing *CCR7* and *ISG15* (**Figure 3A**). These included CCR7⁺ISG15^hi^CADM1⁺ cells that retained the cDC1 lineage marker of *CADM1* and CCR7⁺ISG15^hi^CD1C⁺ cells expressing the cDC2 lineage marker *CD1C* (**Figure 3B**). Trajectory analysis using Monocle3(*32*) suggested that these CCR7⁺ISG15^hi^ populations potentially represent an early state within the maturation pathway of cDC1 and cDC2 transitioning to the CCR7^+^ DC phenotype (**Figure 3C**). Trajectory analysis additionally revealed a maturation pathway originating from the cDC2/DC2-like clusters that converge toward C5_ISG⁺ DC. Given the heterogeneity of the cDC2/DC2-like clusters, we next investigated which cDC2/DC2-like subset might serve as the source of ISG⁺ DCs. Using the transcriptomes of cDC2/DC2-like subtypes as references, SingleR-based stratification(*33*) identified C0_inf-DC2 and C7_inf-DC2 as the most probable origin of C5_ISG⁺ DCs (**Figure 3D and Supplemental Figure 5C**). To further investigate these relationships, we inferred cDC1- and cDC2-derived CCR7^+^ DCs by stratifying C4_CCR7^+^ DCs using the transcriptomes of CCR7⁺ISG15^hi^ cells as references (**Figure 3E and Supplemental Figures 5D, E**). A subset of cDC2-derived CCR7^+^ DCs was inferred to originate from C0_inf-DC2, C5_ISG⁺ DC, and C7_inf-DC2 (**Figure 3F**), further supporting a maturation transition from C5_ISG⁺ DCs to C4_CCR7^+^ DCs. Consistent with these findings, analysis of the mouse dataset revealed a similar pattern. Cells located at the interface between CCR7⁺ DC and cDC1 clusters, as well as between CCR7⁺ DC and ISG⁺ DC clusters, exhibited elevated ISG signature scores (**Supplemental Figure 5F**). Correspondingly, trajectory analysis indicated that cDC2 cells converged toward CCR7⁺ DCs through a C6_ISG⁺ DC state (**Figure 3G**). Together, these analyses suggest that ISG⁺ DCs may transition toward the CCR7⁺ DC phenotype, while the presence of a large ISG⁺ DC cluster indicates that a subset of ISG⁺ DCs may also be maintained as an ISG-associated state within the TME.

**Fig. 3.**
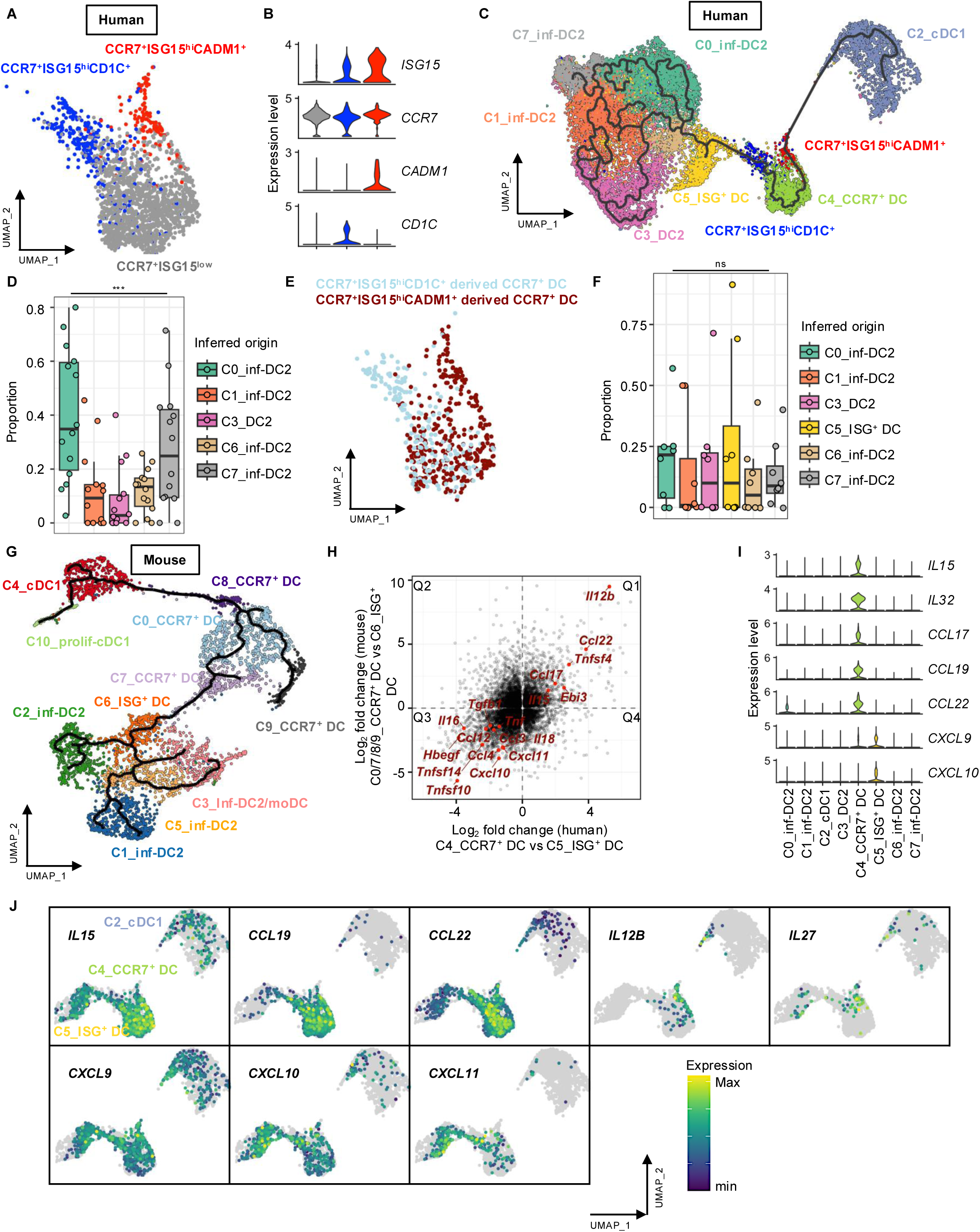
Characterization of activation state of the tumor-infiltrating DCs. **(A)** UMAP analysis of C4_CCR7^+^ DC in human tumor-infiltrating DCs, with colors indicating sub-classification. To identify subpopulations of C4_CCR7^+^ DC, we performed an additional round of integration and re-clustering within C4_CCR7^+^ DC. **(B)** Violin plots showing normalized expression of indicated genes in C4_CCR7^+^ DC sub-clusters, with colors matching those in the UMAP (Figure 3A). **(C)** Monocle 3 analysis of single-cell trajectories for human tumor-infiltrating DCs. Clusters are denoted by colors. **(D)** Boxplot showing the inferred origin fractions of C5_ISG⁺ DC. Dots represent individual samples. Samples lacking any cDC2 subclusters or containing fewer than three C5_ISG⁺ DCs were excluded from the analysis. The lines within the box and whiskers indicate the median and quartiles, respectively. *P*-values (\*\*\**P* < 0.001) were calculated using one-way ANOVA。 **(E)** UMAP analysis of C4_CCR7^+^ DC, with colors indicating inferred cDC1-derived (CCR7^+^ISG15^hi^CADM1^+^) and cDC2-derived (CCR7^+^ISG15^hi^CD1C^+^) DCs. **(F)** Boxplot showing the inferred origin fractions of inferred cDC2-derived CCR7^+^ DCs. Dots represent individual samples. Samples lacking any cDC2 subclusters or containing fewer than two inferred cDC2-derived CCR7^+^ DCs were excluded from the analysis. The lines within the box and whiskers indicate the median and quartiles, respectively. *P*-values were calculated using one-way ANOVA. ns: not significant. **(G)** Monocle 3 analysis of single-cell trajectories for mouse tumor-infiltrating DCs. Clusters are denoted by colors. **(H)** Scatter plots showing log₂ fold changes in gene expression comparing CCR7^+^ DCs vs ISG⁺ DCs in humans and mice. Human genes were mapped to mouse orthologs to enable cross-species visualization. **(I)** Violin plots showing normalized expression of indicated genes in human tumor-infiltrating DCs. **(J)** UMAP displaying normalized gene expression of indicated genes in clusters (C5_ISG⁺ DC, C4_CCR7^+^ DC, and C2_cDC1). Cells with expression levels below 10⁻⁹ are displayed in gray.

### ISG⁺ DC and CCR7^+^ DC Represent Distinct States with Unique Cytokine/Chemokine Profiles

To compare the activation states of ISG⁺ DCs and CCR7⁺ DCs, we examined DEGs distinguishing these two populations that were largely conserved between murine and human datasets (**Figure 3H**). Most of the top 100 up- and downregulated DEGs were located in quadrants 1 and 3 (120 and 148 genes, respectively), whereas quadrants 2 and 4 contained fewer genes (50 and 38), indicating concordant regulation (**Figure 3H**). Notably, several DEGs encoded cytokines and chemokines, prompting us to examine whether these DC activation states differ in their cytokine and chemokine transcriptional profiles. Indeed, these two subsets in humans exhibited distinct cytokine profiles, with C4_CCR7^+^ DC expressing *IL15*, *IL32*, *CCL17*, *CCL19*, and *CCL22*, whereas C5_ISG⁺ DC expressed *CXCL9* and *CXCL10* (**Figures 3I, J, Supplemental Figure 5G, and Supplemental table S7**). Additional heterogeneity in cytokine/chemokine profiles was observed depending on whether the DCs were cDC1- or cDC2-derived. In line with previous studies suggesting that *IL12B* expression is specific to cDC1-derived CCR7^+^ DCs and that IL-12 production depends on IFN-γ signaling in cDC1s(*12, 14*), we found that *IL12B* was specifically expressed in the CCR7⁺ISG15^hi^CADM1⁺ subpopulation (**Figure 3J**). This same subpopulation also showed higher expression of *IL27*, a cytokine that promotes a sustained cytotoxicity CD8 T cell program while limiting their terminal exhaustion(*34*). While *CXCL9* was broadly expressed in C2_cDC1, C4_CCR7^+^ DC, and C5_ISG⁺ DC, *CXCL10* was more highly expressed in C5_ISG⁺ DC and cDC2-derived CCR7^+^ DCs (**Figure 3J and Supplemental table 7**). In mouse datasets, a similar pattern was observed: CCR7⁺ DCs positioned proximal to ISG⁺ DCs or cDC1s in UMAP space exhibited a pronounced IFN signature, with enrichment of *Cxcl9* and *Cxcl10* toward the ISG⁺ DC side and *Il12b* expression toward the cDC1 side (**Supplemental Figure 5F, H**).

### Spatial Distribution of DC Subsets

We then asked whether CCR7^+^ DCs and ISG^+^ DCs differ in their spatial localization within the TME. To address this, we analyzed paired samples of scRNA-seq and spatial transcriptomics (ST) data from three human colorectal cancer (CRC) patients, which included a scRNA-seq reference generated from serial sections of the same formalin-fixed paraffin-embedded (FFPE) blocks(*35*) (**Supplemental Figures 6A–F**). By using scRNA-seq as reference, we performed spot deconvolution to classify and label 8-µm bins with cell types derived from a single cell (**Figure 4A**, **Supplemental Figures 7A–C** and **see methods**). When each DC subtype was highlighted, CCR7^+^ DCs and ISG⁺ DCs displayed a dense and localized distribution pattern, in contrast to the more scattered arrangement of cDC1 and cDC2 (**Supplemental Figures 8A–C**). Although all DC subtypes were present within large lymphoid aggregates in CRC patient1, density mapping revealed distinct aggregation sites for CCR7⁺ DCs and ISG⁺ DCs (**Figures 4B**). CCR7⁺ DCs preferentially localized to regions enriched in *CCL19* and CCL21 (**Figure 4C**), consistent with prior reports showing that CCR7⁺ DCs favorably accumulate within intratumoral niches enriched in CCR7 ligands, produced by cancer-associated fibroblasts (CAFs) and perivascular cells(*36*). In contrast, *CCL19* and *CCL21* expression was not apparent in regions enriched for ISG⁺ DCs (**Figure 4C**). Although CAFs and endothelial cells were broadly distributed throughout the TME, *CCL19*- and *CCL21*-expressing cells were enriched in regions proximal to CCR7⁺ DCs (**Figure 4D**).

**Fig. 4.**
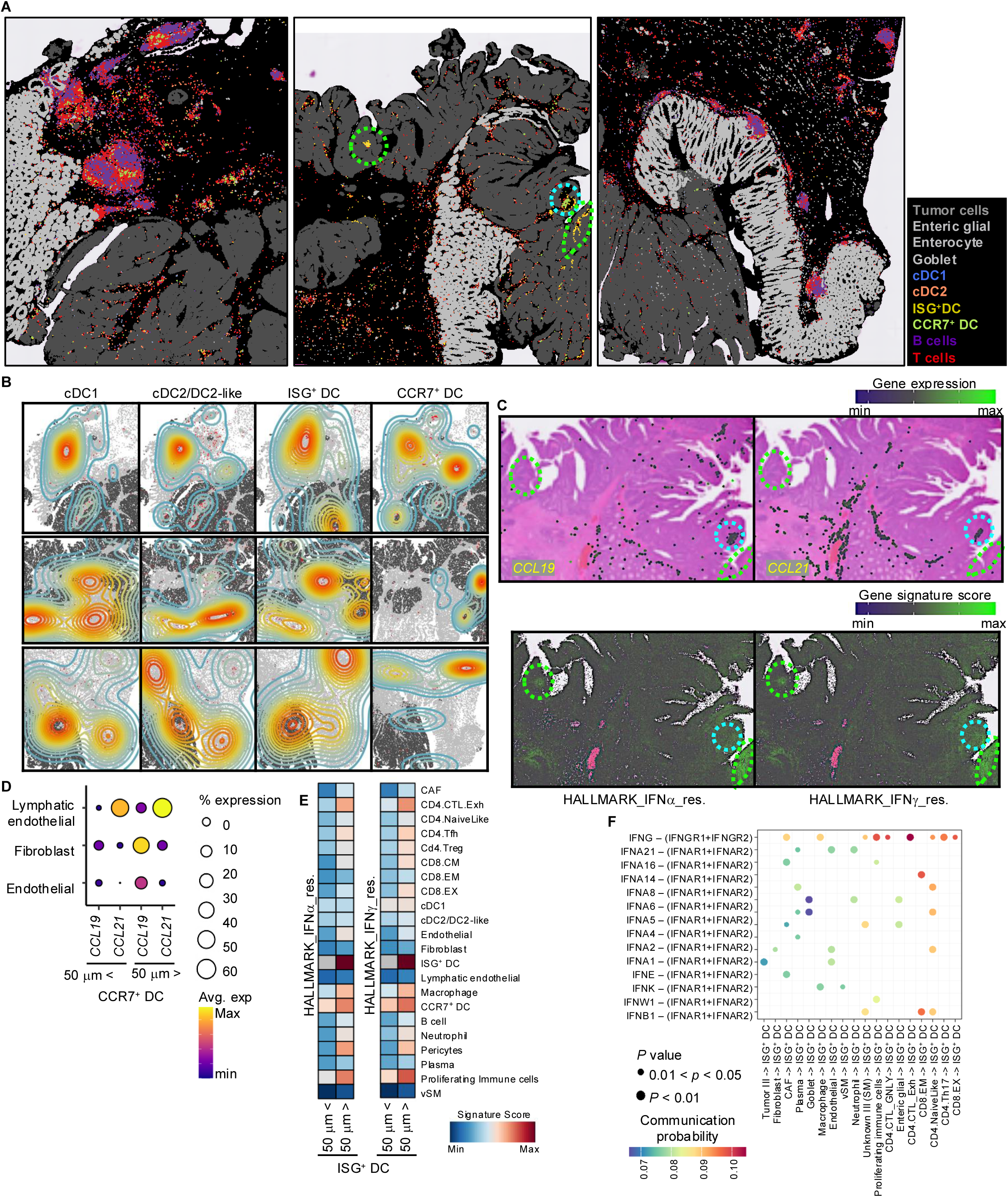
Spatial distribution of DCs in the tumor microenvironment. **(A)** Spatial mapping of three colorectal cancer (CRC) samples (left: CRC patient1, middle: CRC patient2, right: CRC patient5). DC subtypes, B cells, T cells, tumor cells, and enteric glial/enterocyte/goblet cells, predicted by deconvolution using the scRNA-seq reference dataset, are shown. Regions outlined in green and cyan color indicate selected examples of ISG⁺ DC- and CCR7⁺ DC-enriched areas, respectively, which are further shown in Fig. 4C. **(B)** Kernel density maps highlighting the spatial enrichment of DC subtypes (top: CRC patient1, middle: CRC patient2, bottom: CRC patient5). **(C)** Spatial expression of *CCL19* and *CCL21* (top), and IFN-related gene signature scores (bottom) in CRC patient2, overlaid on H&E staining (top). Signature score was calculated using the UCell package(*74*). Regions outlined in green and cyan color indicate ISG⁺ DC- and CCR7⁺ DC-enriched areas, respectively, as shown in Fig. 4A. **(D)** Dot plots illustrating the expression levels of *CCL19* and *CCL21* in CRC patient2, the indicated cells located proximal (< 50 µm) or distal (> 50 µm) to CCR7⁺ DCs. **(E)** Heatmap showing IFN-related gene signature scores across cell types located proximal (< 50 µm) or distal (> 50 µm) to ISG⁺ DCs. **(F)** Bubble plot showing spatial ligand-receptor interactions mediated by type I and type II IFNs from neighboring cell types to ISG⁺ DCs, inferred by CellChat analysis.

In contrast, ISG⁺ DCs were positioned within regions characterized by IFN signaling activity (**Figure 4C**), and multiple neighboring cells within these regions also exhibited elevated IFN signatures (**Figure 4E**). To identify putative cellular sources of IFNs, we performed spatial cell-cell communication analysis using CellChat(*37*). This analysis indicated that multiple immune and non-immune cell types contribute IFN signals to ISG⁺ DCs within these regions (**Figure 4F**). Collectively, these results indicate that CCR7⁺ DC and ISG⁺ DC subsets occupy partially overlapping yet distinct spatial niches within the TME, shaped by region-specific signaling environments.

### Differential Ligand-Receptor Activities in T Cell Enrichment

Given the distinct cytokine and chemokine transcriptional profiles of ISG⁺ DCs and CCR7⁺ DCs, we next asked whether these DC subsets occupy different immune cell neighborhoods within the TME. Neighborhood enrichment analysis indicated that both DC subsets localized within T cell-enriched regions; however, the phenotypic composition of T cells in proximity to each DC subset differed. Specifically, CCR7⁺ DCs exhibited strong enrichment for multiple CD4 T cell phenotypes, including naïve CD4 T cells (CD4. NaïveLike), CD4 T follicular helper (Tfh) cells (CD4.Tfh), CD4 Th17 cells (CD4.Th17), and CD4 T regulatory cells (Tregs) (CD4.Treg) (**Figure 5A, B**), a finding further supported by nearest-neighbor distance analysis (**Figure 5C**). Spatial cell-cell communication analysis suggested that *CCL22* derived from CCR7⁺ DCs may contribute to the recruitment of these CD4 T cell subsets via *CCR4* (**Supplemental Figure 8D**). In contrast, regions enriched for ISG⁺ DCs were characterized by close proximity to CD8 T cells expressing exhaustion-associated transcripts (CD8.EX), potentially encompassing heterogeneous populations, such as activated, effector-like exhausted, and dysfunctional T cells (**Figures 5A–C and see Methods**). Consistent with the prominent IFN signaling observed in these regions, foci of the IFN-inducible chemokines *CXCL9* and *CXCL10* overlapped with ISG⁺ DC-rich areas (**Figure 5D**). Notably, while *CXCL9* and *CXCL10* were also detectable in CCR7⁺ DCs, their expression was markedly more prominent in ISG⁺ DC-enriched regions (**Figure 5D**). In line with the broad IFN signature, multiple immune cell types, including macrophages and T cells, as well as non-immune cells such as endothelial cells and pericytes, expressed these chemokines within ISG⁺ DC regions, potentially establishing these areas as IFN-driven immune hotspots (**Figure 5E**). Spatial cell-cell communication analysis further suggested that *CXCL9* and CXCL*10* produced by ISG⁺ DCs may engage *CXCR3* on neighboring T cells (**Supplemental Figure 8D**). Notably, proliferating immune cells were enriched within both CCR7⁺ DC- and ISG⁺ DC regions (**Figure 5A–C**). Collectively, these findings indicate that CCR7⁺ DC and ISG⁺ DC niches share partially overlapping cytokine and chemokine signaling programs, but differ in the relative contribution and dominance of specific ligand-receptor axes, with certain signaling pathways being preferentially associated with CCR7⁺ DCs. Together, these regionalized signaling differences likely shape distinct yet overlapping T cell-enriched microenvironments within the tumor.

**Fig. 5.**
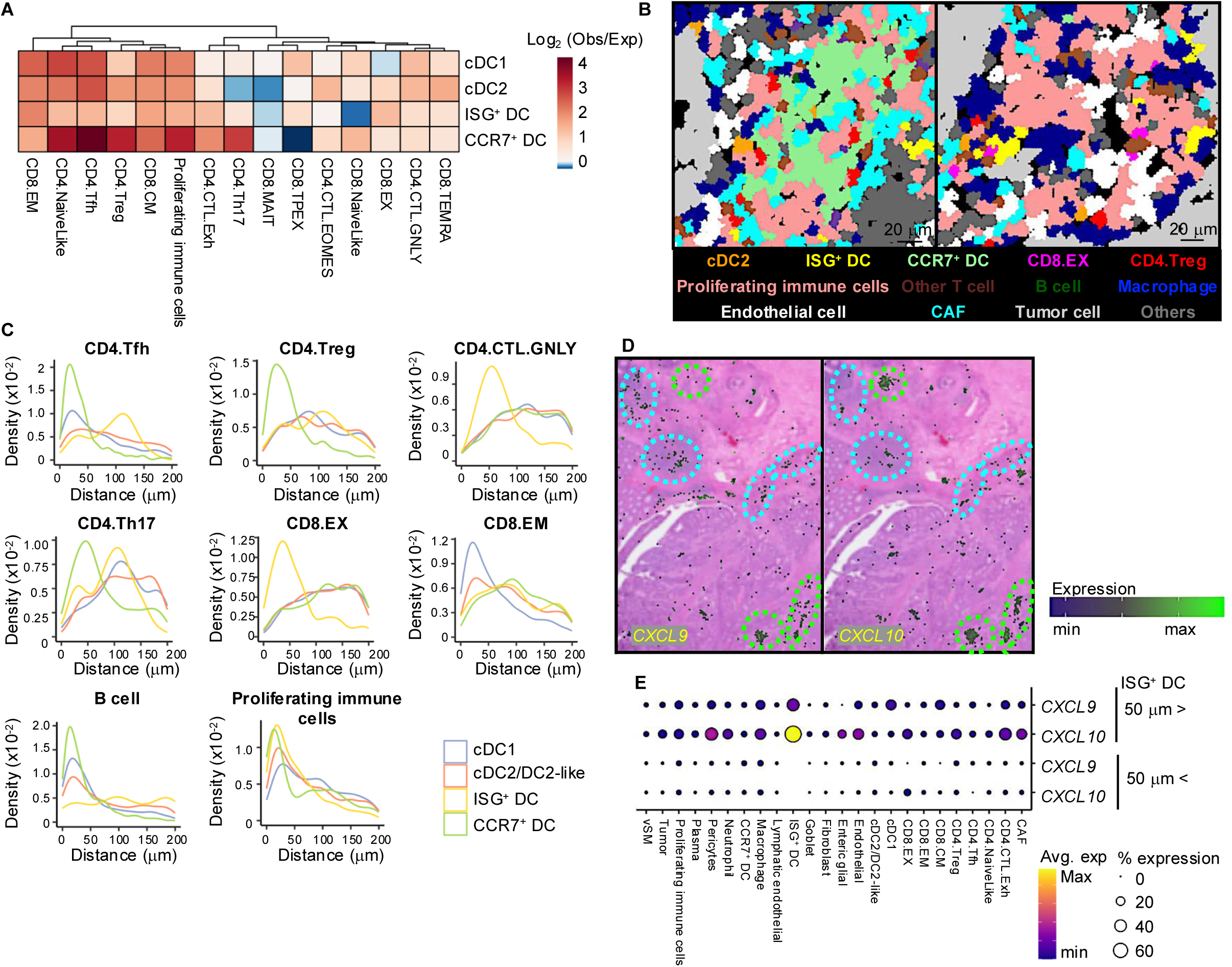
Distinct ligand-receptor axes operating in CCR7^+^ DC and ISG⁺ DC regions. **(A)** Neighborhood enrichment analysis showing the spatial co-occurrence of DC subsets and T cell phenotypes. **(B)** Representative spatial maps of CCR7⁺ DC (left) and ISG⁺ DC (right) enriched regions. Segmented cells are colored by selected annotated cell types. **(C)** Nearest-neighbor distance analysis showing the spatial proximity of each DC subtype to selected immune cell populations **(D)** Spatial expression of *CXCL9* and *CXCL10*, overlaid on H&E staining. Regions outlined in green and cyan indicate ISG⁺ DC- and CCR7⁺ DC-enriched areas, respectively. **(E)** Dot plots illustrating the expression levels of *CXCL9* and CXCL10 in cells proximal (< 50 µm) or distal (> 50 µm) to ISG⁺ DCs.

### Construction of Single-cell Spatial Maps of Immune Cells Across Immunotherapies

Accumulating evidence has demonstrated that spatial organization and physical interaction between DC and T cells are key determinants of effective immunotherapy responses(*9, 10, 38*). Motivated by this, we sought to resolve the spatial architecture of DCs and T cells across distinct immunotherapeutic conditions by generating single-cell-resolved spatial maps using co-detection by indexing (CODEX; also known as PhenoCycler) (*39*) multiplexed tissue imaging (**Figure 6A**). We employed a previously established Y1.7LI murine melanoma model, in which tumor-infiltrating immune cells have been well characterized by scRNA-seq(*40*). This model is responsive to either αPD-1 or αCTLA-4 ICT alone or in combination, as well as a neoantigen synthetic long-peptide (SLP) + polyinosinic-polycytidylic acid (polyI:C) vaccine (NeoVAX), enabling direct comparison of distinct immunotherapy modalities independent of differences in overall therapeutic efficacy. Over 1.4 million cells across 20 samples of whole-tumor sections from Y1.7LI tumor-bearing mice treated with ICTs, NeoVAX, and appropriate controls were analyzed using a custom panel of more than 30 markers, enabling simultaneous detection of a broad range of immune cell phenotypes and functional states (**Supplemental Table S8**, **Supplemental Figure 9A, and see Methods**). To enable precise cell-type annotation, we integrated matched scRNA-seq reference data(*40*) with CODEX protein profiles using the matching cross-modality via fuzzy smoothed embedding (MaxFuse) algorithm(*41*), complemented by manual curation (**Supplemental Figures 9B–D**). This approach identified 10 major immune cell types, with further subdivision of DCs and T cells into distinct subtypes. We validated these annotations using three complementary approaches. First, protein expression patterns detected by CODEX were concordant with corresponding gene expression signatures observed in scRNA-seq data across cell types (**Figure 6B, and Supplemental Figure 9E**). Second, relative cell-type proportions were consistent between scRNA-seq and CODEX datasets (**Supplemental Figure 9F**). Third, the spatial distribution of annotated cell populations aligned with tissue-level staining patterns of established lineage markers (**Figure 6C**).

**Fig. 6.**
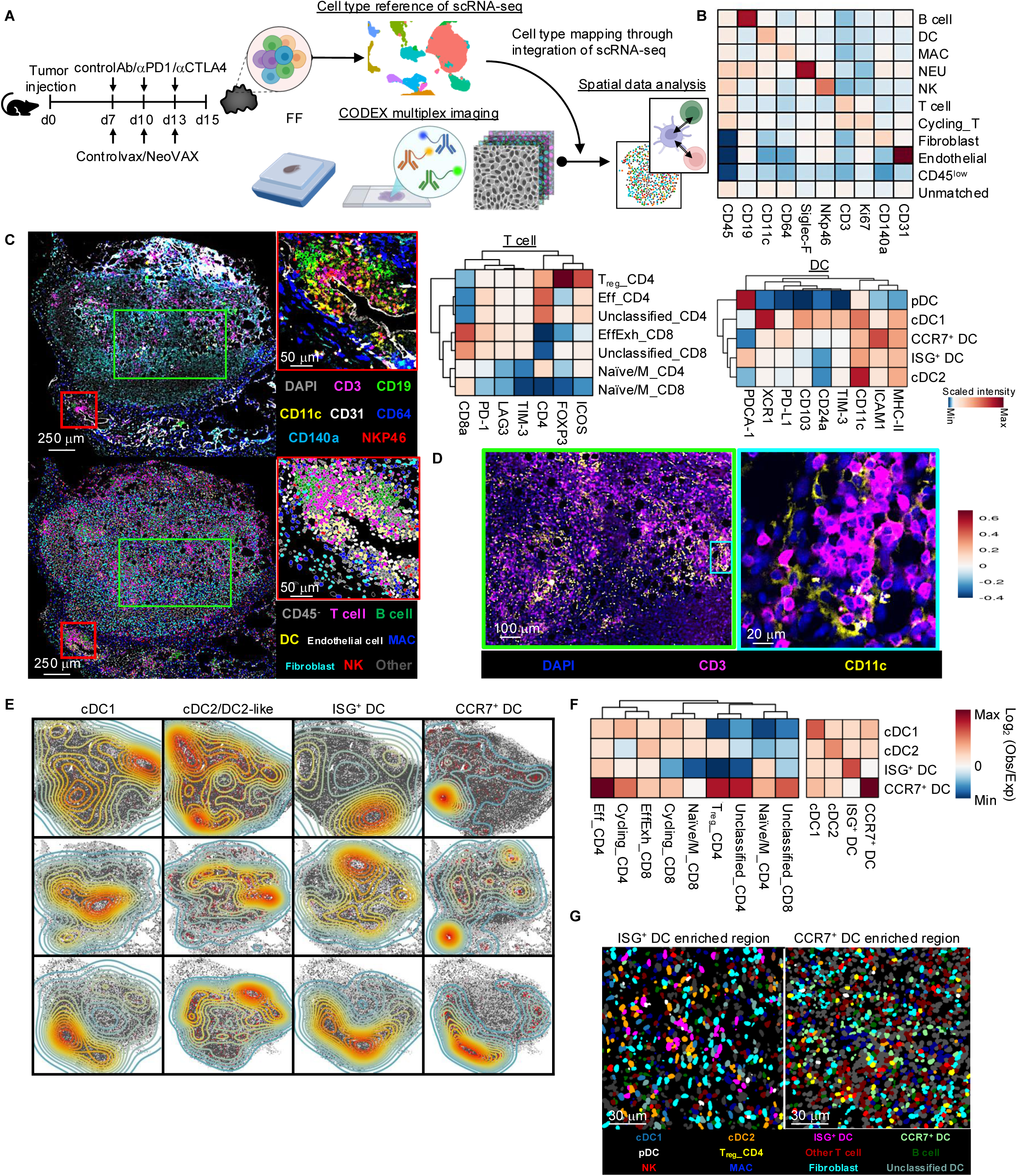
Multiplex imaging profiling of DC subsets and T cell phenotype in a murine tumor model. **(A)** Schematic overview of the experimental and analytical workflow. Tumor-bearing mice were treated with αPD-1, αCTLA-4, αPD-1 and αCTLA-4 combination therapy, neoantigen synthetic long-peptide [SLP] + polyI:C vaccination (NeoVAX), or corresponding control conditions (control monoclonal antibody [control Ab] and irrelevant SLP + polyI:C [control vax]) (six treatment groups; three independent experiments; total n = 20 tumor sections). Fresh frozen tumor sections collected on day 15 were subjected to CODEX multiplexed tissue imaging, and the resulting data were integrated with scRNA-seq reference datasets from the same tumor model to enable cell-type annotation and spatial data analysis. **(B)** Heatmap showing normalized protein staining intensities for scRNA-seq-derived cell type annotations following cross-modality integration. **(C)** Representative CODEX multiplexed imaging of tumor section treated with a control antibody, showing staining patterns of canonical markers (top) and cell segmentation colored by cell-type annotations (bottom). **(D)** Representative CODEX images highlighting the spatial co-localization of T cells and DCs in tumor treated with a control antibody. **(E)** Kernel density maps highlighting the spatial enrichment of DC subtypes in tumors treated with a control antibody across three independent experiments. Individual DC cells are shown as red dots. **(F)** Neighborhood enrichment analysis showing the spatial co-occurrence of DC subsets and T cell phenotypes. **(G)** Representative spatial maps of ISG⁺ DC enriched and CCR7⁺ DC enriched regions. Segmented cells are colored according to selected annotated cell types, while all other cell types are shown in grey.

We next examined the spatial localization of DCs and T cells within the TME of control Ab-treated progressively growing tumors. As expected, DCs were concentrated within T cell-enriched regions (**Figure 6D**). Density mapping and neighborhood enrichment analyses revealed that ISG⁺ DCs and CCR7⁺ DCs localize partially overlapping yet distinct spatial niches and preferentially associate with different T cell subsets, consistent with spatial patterns observed in human tumors (**Figures 6E, F**). Notably, CCR7⁺ DCs displayed spatial proximity to CD4 Tregs (**Figures 6F, G**), consistent with previously described mregDCs-Treg niches(*42*), supporting the robustness of our spatial profiling approach. Collectively, these analyses establish a single-cell spatial framework for interrogating immunotherapy-dependent organization of DCs and T cells within the TME.

### Immunotherapy-Induced Remodeling of CCR7⁺DCs and ISG⁺DCs Hubs

Using this spatial dataset, we next focused on immunotherapy-induced remodeling of immune hubs formed by CCR7⁺ DCs and ISG⁺ DCs within the TME. Cellular neighborhood analysis revealed a pronounced increase in homotypic enrichment of CCR7⁺ DCs specifically in αPD-1-treated tumors (**Figure 7A**). Consistent with this finding, nearest-neighbor distance analysis demonstrated a marked increase in the density of CCR7⁺ DCs at shorter intercellular distances, indicating enhanced spatial clustering in αPD-1-treated tumors (**Figure 7B**). In line with these observations, CCR7⁺ DCs formed larger spatial clusters in αPD-1-treated tumors compared with control Ab and exhibited increased proximity to EffExh_CD8 and cycling_CD8 (**Figures 7C–E and Supplemental Figures 10A, B**). A similar trend toward enhanced homotypic clustering was also observed for ISG⁺ DCs in αPD-1-treated tumors (**Figures 7A, B, D and Supplemental Figure 10C**). In addition, ISG⁺ DCs exhibited increased spatial proximity to EffExh_CD8 and Eff_CD4 under αPD-1 treatment compared with control Ab (**Figure 7E**). Given previous evidence that effective αPD-1 therapy is partially driven by local expansion and effector differentiation of intratumoral CD8 T cells within DCs and T helper-rich niches(*38*), the observed spatial rearrangements suggest a potential contribution of CCR7⁺ DC and ISG⁺ DC hubs to antitumor immune activity. Notably, scRNA-seq analyses of human and mouse tumor-infiltrating DCs, as well as of the corresponding tumor model, revealed minimal alterations in DC composition or transcriptional states across treatment conditions (**Supplemental Figures 11A–F**).

**Fig. 7.**
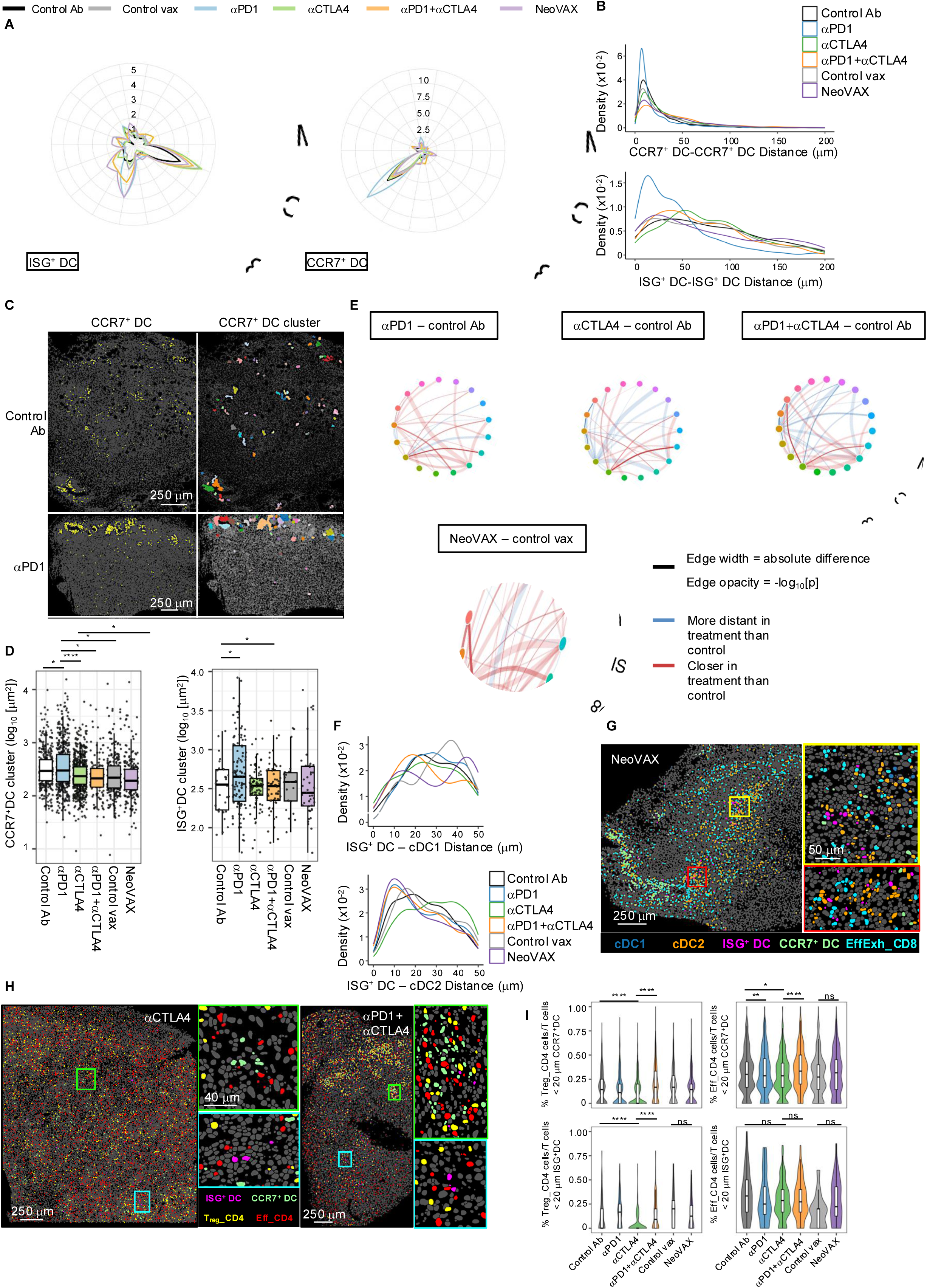
Immunotherapy-dependent remodeling of CCR7⁺ DC and ISG⁺ DC spatial hubs. **(A)** Radar plots summarizing neighborhood enrichment profiles of ISG⁺ DCs (left) and CCR7⁺ DCs (right) across treatment conditions. Values represent the ratio of observed to expected enrichment, indicating preferential spatial association or avoidance between cell types. **(B)** Nearest-neighbor distance analysis showing the spatial proximity of homotypic DC pairs: CCR7⁺ DC-CCR7⁺ DC distances (top) and ISG⁺ DC-ISG⁺ DC distances (bottom) across treatment conditions. **(C)** Representative spatial maps illustrating CCR7⁺ DC distribution (left) and CCR7⁺ DC clusters (right) under control antibody and αPD-1 treatment conditions. Clusters were defined using Density-Based Spatial Clustering of Applications with Noise (DBSCAN). **(D)** Boxplots showing the size of CCR7⁺ DC cluster area (left) and ISG⁺ DC cluster area (right) across treatment conditions. Each dot represents the individual cluster. The lines within the box and whiskers indicate the median and quartiles, respectively. P-values (*P < 0.05, **P < 0.01, ***P < 0.001, ****P < 0.0001) were calculated using one-way ANOVA with Tukey’s multiple comparison test. ns: not significant. **(E)** Chord diagrams of distance permutation analysis, performed by SPACEc, depicting treatment-induced changes in spatial neighborhood distances relative to respective control treatment conditions. **(F)** Nearest-neighbor distance analysis showing the spatial proximity of ISG⁺ DCs to cDC1s (top) and to cDC2s (bottom) across treatment conditions. **(G)** Representative spatial maps from NeoVAX-treated tumors illustrating the proximity of ISG⁺ DCs to cDC1s, cDC2s, and EffExh_CD8 T cells. **(H)** Representative spatial maps showing the distribution of Eff_CD4 and Treg_CD4 T cells around CCR7⁺ DCs and ISG⁺ DCs under αCTLA-4 treatment (left) and combination therapy (right). **(I)** Violin plots depicting the proportion of Eff_CD4 and Treg_CD4 T cells within 20 µm of CCR7⁺ DCs and ISG⁺ DCs. P-values (*P < 0.05, **P < 0.01, ***P < 0.001, ****P < 0.0001) were calculated using one-way ANOVA with Tukey’s multiple comparison test. ns: not significant.

Beyond αPD-1-specific effects, increased enrichment between cDC2 and ISG⁺ DCs was observed across ICT conditions, with an even more pronounced effect detected following NeoVAX treatment (**Figures 7A, F**). In the same tumor model, we previously demonstrated that NeoVAX robustly induces neoantigen-specific CD8 T cells, including proliferating, stem-like progenitor exhausted, as well as IFN-γ^high^CD8 T cells(*40*). Given that cDC2-ISG⁺ DC neighborhoods are located within regions enriched for EffExh_CD8 T cells (**Figure 7G**), these spatial features together raise the possibility that local IFN-γ signaling from neoantigen-reactive CD8 T cells promotes the transition of cDC2s toward an ISG⁺ DC state within the tumor.

### Combination αCTLA-4 and αPD-1 ICT Exhibits Non-Additive Effects in CCR7⁺ DC and ISG⁺ DC Hubs

The combination of αCTLA-4 and αPD-1 therapy did not uniformly amplify the potentially favorable spatial features observed under monotherapy. Instead, the combination of αCTLA-4 and αPD-1 therapy both enhanced and attenuated spatial configurations observed under monotherapy. One attenuating effect was the absence of CCR7⁺ DC cluster expansion under combination ICT relative to αPD-1 (**Figures 7C, D** and **Supplemental Figure 10B**). Another attenuating effect was the enrichment of Tregs. Whereas αCTLA-4 monotherapy reduced the overall abundance of Tregs, consistent with the known activity of the αCTLA-4 clone 9D9(*43*), as well as Tregs in the vicinity of both CCR7⁺ DCs and ISG⁺ DCs (**Figures 7H, I**), combination therapy instead resulted in increased Treg accumulation around both CCR7⁺ DCs and ISG⁺ DCs compared with αCTLA-4 monotherapy, with a particularly pronounced enrichment around CCR7⁺ DCs (**Figure 7I and Supplemental Figure 10A**).

Regarding enhancing effects, we and others have previously observed that αCTLA-4 treatment increases ICOS⁺PD-1⁺ Th1 CD4 T cells within the TME(*40, 44–46*), and that combination therapy further promotes the expansion of Th1 CD4 T cells(*40, 47*). Under αCTLA-4 monotherapy, spatial co-localization between Eff_CD4, a population that includes Th1 cells, and DCs was not increased (**Figure 7I and Supplemental Figure 10A**). In contrast, combination therapy induced increased enrichment of Eff_CD4 cells in proximity to CCR7⁺ DCs (**Figure 7I and Supplemental Figure 17A**). This spatial reorganization suggests that enhanced local DC-CD4 T cell interactions may contribute to the Th1 expansion in the context of combination therapy.

Collectively, these spatial analyses show that CCR7⁺ DC and ISG⁺ DC hubs undergo immunotherapy-dependent remodeling and are associated with the spatial redistribution of T cell subsets, suggesting that both DC states play a role in immunotherapy-induced T cell pool within the TME.

## DISCUSSION

In this study, we constructed an integrative atlas of tumor-infiltrating DCs across human and murine tumors spanning multiple cancer types and treatment conditions. By harmonizing scRNA-seq with spatial transcriptomics and multiplexed imaging, we delineated conserved DC transcriptional programs together with their spatial organization in the TME. Although CCR7⁺ DC and ISG⁺ DC states have been described previously, our study extends these observations by demonstrating that these states form spatially organized immune hubs embedded within distinct signaling niches, associate with different neighboring T cell phenotypes, and undergo immunotherapy-dependent remodeling.

One of the key findings from transcriptome analysis is a refined view of the ISG⁺ DC population and its conservation across cancer types and species. Our analysis identified three DC populations exhibiting high ISG signature. Among them, CCR7^+^ISG15^hi^CADM1^+^ and CCR7^+^ISG15^hi^CD1C^+^ subpopulations within the CCR7⁺ DC compartment potentially represented transient ISG activation in cDC1-derived and cDC2-derived CCR7^+^ DCs, respectively. This finding aligns with previous studies suggesting that both cDC1s and cDC2s can transiently engage ISG transcriptional programs(*3, 20, 21*). Importantly, our data extend this concept by revealing a substantial population of ISG⁺ DCs aligned with the cDC2 lineage (C5_ISG⁺ DC), which is distinct from CCR7⁺ DCs not only at transcriptional level but also at spatial organization. This observation raises a fundamental question of lineage and plasticity, whether ISG⁺ DCs constitute a transient activation intermediate preceding CCR7⁺ DC differentiation, or instead represent a stable, functionally maintained state within the TME. Our analyses are compatible with both possibilities. Developmental inference, together with the partial spatial co-localization observed between ISG⁺ DCs and CCR7⁺ DCs, supports a model in which cDC2/DC2-like states transition toward ISG⁺ DCs and subsequently differentiate into CCR7⁺ DC phenotypes. On the other hand, the presence of a sizable ISG⁺ DC compartment and its localization within spatial niches separated from CCR7⁺ DC hubs suggest that at least a fraction of ISG⁺ DCs may persist as a stable, spatially constrained state rather than a mere transient intermediate.

Spatial transcriptomics analysis provides insight into how these DC activation states (CCR7^+^ DCs and ISG⁺ DCs) are organized within the TME. As anticipated, CCR7^+^ DCs preferentially localized to stromal niches enriched for CCR7 ligands where they form DC-T cell immune hubs. ISG⁺ DCs resided in distinct niches characterized by a strong IFN signature and focal enrichment of IFN-inducible chemokines, particularly CXCL9 and CXCL10. Notably, although CXCL9/10 could be detected in multiple immune contexts, their spatial enrichment was most pronounced in the ISG⁺ DC regions. This observation also raised questions concerning the upstream factors that induce and sustain ISG⁺ DC hubs. Our spatial communication analyses further suggest that IFN signals driving ISG⁺ DC states may originate from diverse immune and non-immune cellular sources within IFN-active regions, including CD4 T cells, CD8 T cells, tumor cells and CAFs. This distributed signaling suggests a model in which ISG⁺ DC hubs emerge as a consequence of localized inflammatory circuits shaped by convergent IFN input from multiple cell types rather than a single dominant cellular driver. Such circuits may reinforce ISG programs in DC while simultaneously shaping chemokine landscapes that influence T cell recruitment and positioning.

We leveraged a well-characterized murine melanoma model together with large-scale, single-cell-resolved spatial imaging. Using CODEX with a panel of more than 30 markers, we generated high-dimensional spatial maps across multiple immunotherapeutic conditions, including αPD-1, αCTLA-4, αPD-1 plus αCTLA-4, and NeoVAX, alongside two control settings in which tumors progress. Importantly, we integrated these spatial datasets with matched scRNA-seq profiles generated from the same tumor model and treatment conditions. This experimental design enabled direct, within-model comparison of immunotherapy-induced remodeling of DC-T cell spatial niches, providing a level of multimodal and therapeutic resolution that has been challenging to achieve in prior studies.

Multiplexed imaging analyses demonstrate that αPD-1 enhances homotypic clustering of CCR7⁺ DCs and, to a lesser extent, ISG⁺ DCs, accompanied by increased spatial proximity to effector and cycling CD8 T cells. This supports the idea that effective therapy amplifies hub-like microenvironments coupled with local T cell expansion. Moreover, the enhanced enrichement of cDC2-ISG⁺ DC neighborhood, following NeoVAX treatment, suggests a potential feedback loop in which neoantigen-specific IFN-γ^high^ CD8 T cells promote IFN-exposed DC. At the same time, our analyses highlight that combination therapy does not simply represent an additive intensification of favorable spatial features. The expansion of intratumoral CCR7⁺ DC clusters in response to αPD1 was not recapitulated under combination therapy, which may reflect a qualitatively distinct remodeling of CCR7⁺ DC. Building on recent work demonstrating that Tregs establish peri-lymphatic niches that restrict mregDC migration to tdLNs(*42*), our data suggest that αCTLA-4 may relieve this spatial constraint, facilitating CCR7⁺ DC migration to tdLNs, thereby limiting the persistence of large intratumoral clusters. The functional consequences of this alternative remodeling for anti-tumor immunity remain to be elucidated.

ISG⁺ DCs have been reported to promote anti-tumor CD8 T cell immunity and tumor rejection even in *Batf3*^−/−^ mice lacking cDC1(*23*). In this context, the enrichment of proliferating CD8 T cells and CD8 T EX cells within ISG⁺ DC regions in human tissue is particularly relevant. While transcriptional exhaustion is frequently observed among tumor-specific CD8 T cells(*48, 49*), exhaustion does not uniformly equate to loss of effector function. Rather, accumulating evidence supports a spectrum of exhaustion states characterized by progressive functional impairment, ranging from early exhausted cells that retain substantial effector capacity to terminally exhausted cells with more pronounced dysfunction(*50*). In this framework, stem-like CD8 T cells represent a progenitor population that sustains the exhausted pool but is itself not functionally exhausted. The co-occurrence of proliferating CD8 T cells and enhanced IFN signaling within ISG⁺ DC regions therefore suggests that these niches may support functional, tumor-specific T cells responses despite the presence of exhaustion-associated transcriptional programs.

Our study has several limitations. First, much of our analyses were based on transcriptomic data, and further validation at the protein and functional levels is necessary. Second, trajectory analyses provide computational predictions of cellular state relationships but do not constitute direct lineage tracing or imply sequential developmental stages. Finally, our spatial transcriptomics analyses in human samples were limited in cohort size, and additional patient datasets across tumor types will be needed to establish the generality of specific niche architectures.

In conclusion, our comprehensive study delineates the phenotypic heterogeneity of tumor-infiltrating DCs alongside their spatial distribution and co-localization with T cells. Understanding how immunotherapies remodel DC-T cell niches may provide insights into potential strategies to enhance immunotherapeutic efficacy by modulating the local DC landscape. Beyond these biological insights into DC immunology, our DC atlas also serves as a resource with broad applicability for future studies of tumor immunity and immunotherapy.

## MATERIALS AND METHODS

### Mice

Wildtype (WT) C57BL/6J mice were purchased from The Jackson Laboratory (JAX, Bar Harbor, ME, USA). In vivo experiments were performed using 8- to 12-week-old male mice. Mice were housed in a specific pathogen-free animal facility. All animal studies were conducted in accordance with protocol and the approval of the Institutional Animal Care and Use Committee (IACUC) of The University of Texas MD Anderson Cancer Center (Houston, TX).

### Tumor cell line

Y1.7LI(*40*) is a murine melanoma cell line that we previously generated from the *Braf^V600E^Pten*^−/−^*Cdkn2a*^−/−^ YUMM1.7 line background by engineering expression of defined MHC class I (*Lama4*^G1254V^) and class II (*Itgbl*^N710Y^) neoantigens(*51, 52*).

### Tumor transplantation

Y1.7LI melanoma cells were cultured in R-10 plus BME medium consisting of RPMI-1640 (HyClone) supplemented with 1% L-glutamine, 1% penicillin-streptomycin, 1% sodium pyruvate, 0.5% sodium bicarbonate, 0.1% 2-mercaptoethanol, and 10% heat-inactivated fetal bovine serum (FBS; HyClone). Cells were passaged 3-6 times after thawing before use in experiments. Prior to injection, tumor cells were washed extensively and resuspended at a concentration of 3.33 × 10⁶ cells/mL in endotoxin-free phosphate-buffered saline (PBS). A total volume of 150 μL (5 × 10⁵ cells) was injected subcutaneously into the flank of recipient mice. Cell viability at the time of injection exceeded 90%, as assessed by trypan blue exclusion. Tumor growth was monitored by caliper measurements and expressed as the mean of two perpendicular diameters. Survival endpoints were defined as mouse death or a mean tumor diameter of 20 mm.

### In vivo vaccination, antibody treatments, and tumor tissue processing for CODEX imaging

For ICT treatment, Y1.7LI tumor-bearing mice were treated intraperitoneally with 200 μg of αCTLA-4 and/or αPD-1 on day 7, 10, and 13 post-tumor transplant. For control groups, mice were injected with 200 μg of IgG2a isotype control antibodies (control Ab). For in vivo experiments, “*In vivo* Platinum”-grade antibodies that were verified to be free of mouse pathogens (IDEXX IMPACT I mouse pathogen testing) were purchased from Leinco Technologies: αPD-1 (rat IgG2a clone RMP1-14), αCTLA-4 (murine IgG2b clone 9D9). Y1.7LI tumor bearing male mice were vaccinated subcutaneously with 10 μg of mLama4 (NeoVAX) or mAlg8 SLP (control vax) in combination with 50 μg of VacciGrade™ high molecular weight polyI:C (InvivoGen) diluted in endotoxin-free sterile PBS to a total volume of 150 μL on day 7 and 13 post-tumor transplant. For SLP, peptide sequence used for mLama4; QKISFFDGFEVGFNFRTLQPNGLLFYYT, for mAlg8; AVGITYTWTRLYASVLTGSLV. mLama4 SLP (NeoVAX) served as a relevant SLP and mAlg8 SLP (control vax) served as irrelevant SLP for the Y1.7LI line. Tumors were harvested on day 15, embedded in optimal cutting temperature (O.C.T.) compound (Tissue-Tek), and snap-frozen using liquid nitrogen vapor to preserve tissue architecture. Poly-L-lysine-coated coverslips (Sigma) were prepared according to the manufacturer’s protocol (Akoya Biosciences). Tumor sections were cut at 8 µm thickness using a cryomicrotome and mounted onto poly-L-lysine-coated coverslips.

### CODEX Antibody Panel Design and Optimization

CODEX (also known as PhenoCycler) multiplexed imaging was performed according to the established CODEX staining and imaging protocol(*39*). Imaging was conducted through iterative cycles of annealing and stripping of fluorophore-labeled oligonucleotide barcodes complementary to antibody-conjugated barcodes. The antibody panels used in this study are detailed in Supplemental Table S8. Each antibody was conjugated to a unique oligonucleotide barcode and individually tested and titrated to optimize signal-to-noise ratios. All conjugates were then combined and evaluated in a single multicycle CODEX experiment, and optimal antibody dilution, exposure time, and imaging cycle were determined for each marker.

### CODEX Imaging

Tissue arrays/sections were stained with validated CODEX antibody panels and processed on the CODEX/PhenoCycler instrument (Akoya Biosciences), followed by imaging on a fluorescence microscope (BZ-X series, Keyence Corporation of America). CODEX multiplexed imaging was performed through iterative cycles of stripping, annealing, and imaging of fluorescently labeled oligonucleotides complementary to antibody-conjugated barcodes. All tissue arrays underwent CODEX imaging; run-specific metadata are provided in Supplemental Table S8.

### CODEX data Quality control and processing

The raw TIFF imaging outputs were first examined using QuPath software (https://qupath.github.io/) to assess staining quality and signal-to-noise ratios. Samples exhibiting aberrant marker expression patterns or consistently low signal-to-noise ratios were excluded from further analysis. Image processing, quality control, and segmentation were performed using the Python package SPACEc(*53*) (**Supplemental Figure 9A**). Tile-associated shading artifacts were corrected using a VISTAmap(*54*)-inspired approach. Briefly, tissue regions were defined using the DAPI channel, large-scale background variation was removed, and residual tile-related intensity inhomogeneity was corrected through local baseline estimation and seam smoothing across tiles. Shading correction was applied independently to each marker channel. Cell segmentation was performed using MESMER, a pre-trained deep learning algorithm(*55*), with DAPI used to identify nuclei, and CD45 used to define whole-cell boundaries specifically for immune cells. To refine single-cell fluorescence measurements in densely packed tissues, fluorescence spillover compensation was performed as previously described(*56, 57*), using the implementation provided in SPACEc. Cells in the bottom 1% by area, top 0.1% by area, or bottom 1% by DAPI intensity were excluded to remove segmentation artifacts. To minimize the influence of extreme high intensity outliers, which typically results from non-specific fluorescent artifacts or debris, each marker was subjected to one-sided Winsorization at the upper 99th percentile. Marker intensities were then normalized on a per-image basis using z-score normalization implemented in SPACEc. After normalization, cells exhibiting abnormally high expression across multiple antibodies (defined as the top 2% of summed z-scored intensities and positivity for more than 30 antibodies) were additionally removed. Each image was processed independently. Processed image data were integrated across samples using shared markers via Harmony(*58*).

### CODEX cell annotation

We previously characterized immune cell populations from the same murine tumor model under identical treatment conditions using scRNA-seq(*40*). To obtain high-quality cell-type annotations for the CODEX dataset, we integrated the existing scRNA-seq data with CODEX multiplexed imaging data using MaxFuse(*41*) (**Supplemental Figure 9B**). We refined the previously defined scRNA-seq clustering to facilitate robust validation of annotated cells using CODEX protein markers. First, low-quality clusters and tiny populations, including neutrophils and contaminating fibroblast clusters, were removed from the scRNA-seq dataset because these populations were present at very low frequencies and therefore did not reliably represent the corresponding cell populations observed in the CODEX dataset. Second, although T cells had previously been subdivided into 21 clusters, we reduced the granularity to six major subphenotypes (naive/M_CD4, naive/M_CD8, Eff_CD4, Treg_CD4, EffExh_CD8, and Cycling_T). Finally, DC labels were transferred from the integrated scRNA-seq dataset to this reference. For MaxFuse integration, CODEX cells with CD45 z-scored intensity greater than −0.5 were selected, and cell-type annotation was performed in two stages. In the first round, broad immune lineages were assigned, including DCs, B cells, T cells, cycling T cells, macrophages, and natural killer (NK) cells (**Supplemental Figure 9B**). In the second round, finer subtype annotation was performed separately for T cells and DCs using lineage-specific scRNA-seq reference datasets. Because IgM is a secreted protein that can bind to Fcγ receptor expressing cells, IgM was excluded from the CODEX marker set during integration. Cells that were matched with sufficient confidence (MaxFuse score ≥ 0.4) were assigned the corresponding cell-type labels, whereas cells failing to meet this threshold were designated as unassigned cells. Because we observed a substantial number of cells annotated as B cells despite lacking canonical B cell markers, only CD19 positive cells within the initially annotated B cell population were retained as B cells. For CODEX-imaging specific annotations, Endothelial cells, fibroblasts, and neutrophils were annotated manually based on unsupervised clustering and canonical marker expression (CD31, CD140a, and Siglec-F) (**Supplemental Figure 9C**). Each image was processed independently.

### Neighborhood enrichment analysis

For each cell, the neighboring cells within 20 μm were identified based on Euclidean distance in physical space. To verify that the observed cell-cell adjacency patterns were not due to random chance, we employed a permutation-based statistical test. We first tabulated the frequency of interactions between all cell type pairs as the “observed” count, then generated a null model by repeatedly (1,000 times) shuffling the cell type labels while keeping the spatial position of each cell fixed. This process created a distribution of “expected” interaction frequencies that would occur under a random spatial arrangement. From the observed and expected values, we calculated log₂(observed/expected) ratios and associated false discovery rates (FDR) for each cell type pair.

### Processing of collected single-cell RNA-seq datasets

The scRNA-seq data were collected from previously published datasets (**Supplemental Tables S1 and S4**). Prior to integration, each dataset was processed separately in quality control and cell-type annotation. Quality control was conducted by filtering cells based on mitochondrial content, unique molecular identifiers (UMIs), number of features, and the number of genes per UMI. The filtered expression matrices were normalized using Seurat’s “NormalizeData” function. Highly variable genes (n = 2000) were identified using the “FindVariableFeatures” function with default parameters. For cell-type annotation, we employed the automated cell-type identification tool ScType(*59*). To ensure we did not inappropriately omit any DCs, we extracted not only cells annotated as myeloid DCs, but also those labeled as macrophages, monocytes, and ISG-expressing immune cells. Subsequently, a unified dataset comprising 276,163 cells (human) or 101,161 cells (mouse) was generated through the integration of individual studies (**Supplemental Figures 1A and 3A**). This integration was performed using the batch effect correction algorithm implemented in the Seurat V5 pipeline(*60*), employing anchor-based reciprocal principal component analysis (RPCA). Unsupervised graph-based clustering was then performed on the integrated data. Clusters expressing canonical DC markers (*FLT3*, *ZBTB46*, and *ITGAX*) were identified subsetted, and subjected for downstream analysis. Low quality and putative non-DC contaminants, including clusters expressing T cell and B cell marker genes, were excluded prior to further characterization. A second round of integration was subsequently conducted for the refined DC population using the same approach. The integrated DC dataset was scaled using Seurat’s “ScaleData” function. Principal component analysis (PCA) was performed using the “RunPCA” function. Dimensionality reduction for visualization was carried out using UMAP, followed by clustering with the “FindNeighbors” and “FindClusters” functions. To determine the optimal clustering granularity in an unbiased manner, the clustree package)(*61*) was used to assess the SC3 stability metric across a range of clustering resolutions. SC3 stability quantifies how consistently cell groupings are preserved across resolutions, providing a measure of clustering robustness(*62*). The final clustering resolution was selected based on a local maximum in SC3 stability that maintained higher clustering granularity. To evaluate the reproducibility of heterogenous cDC2 clusters, DC integration was additionally performed using an alternative integration methods. Specifically, Harmony(*58*), FastMNN(*63*), and scVI(*64*) integrationg was tested using the IntegrateLayers function implemented in Seurat.

### Evaluating the heterogeneity within single-cell populations

To evaluate the degree of heterogeneity across major DC subsets, we employed ROGUE(*30*), a universal entropy-based metric designed to quantify single-cell population purity. The ROGUE score ranges from 0 to 1, where a value of 1 indicates a highly homogeneous population lacking distinct gene expression patterns, while 0 reflects maximal heterogeneity.

### Identification of cluster-specific genes

Cluster-specific markers were identified using the limma package (version 3.60.0)(*65*), as implemented in Seurat’s “FindAllMarkers” function with a two-sided nonparametric Wilcoxon rank-sum test and Bonferroni correction. Genes with an adjusted *P*-value less than 0.05 were considered DEGs. For the heatmap representations in Figures 1F and Supplemental Figure 3J, the mean expression of marker genes defined as those expressed in more than 50% of cells within each cluster and ranked among the top 30 by log₂ fold change was used.

### Analysis of gene signature score

Among the DEGs, genes expressed in more than 30% of cells within each cluster were further filtered, and the top 50 genes ranked by log₂ fold change were selected as gene signatures. Individual cells were scored for gene-signature enrichment using Seurat’s “AddModuleScore” function. Signature scores were then averaged across cells within each cluster and cancer type, and the resulting mean scores were used for visualization.

### Analysis of cluster similarity across species

To assess the similarity of DC clusters between mouse and human datasets, we performed a cross-species gene signature analysis. Cluster-specific gene signatures for each cancer type were defined using the same criteria described above (serving as the reference signatures). Human gene symbols were then converted to their corresponding mouse orthologs, and gene signature scores were calculated using the “AddModuleScore” function, applying the human-derived gene sets to the mouse dataset. The average signature score for each cluster within the mouse cohorts was computed, and the scaled scores were used for visualization.

### Trajectory inference

Trajectory analysis was performed using the Monocle3 package (version 1.3.7)(*66*). The Seurat object was converted into the Monocle3 object using the “as.cell_data_set” function. To retain existing clustering and dimensional reduction results from Seurat object, partition labels, cluster identities, and UMAP coordinates were manually transferred into the Monocle3 object. The DC differentiation trajectory was inferred with the default parameters.

### Gene set enrichment analysis

Gene set enrichment analysis was performed using the fgsea package (version 1.30.0) to assess the enrichment of Hallmark gene sets from the Molecular Signatures Database (MSigDB). Hallmark gene sets were obtained using the msigdbr package (version 7.5.1). Differential gene expression statistics were calculated using the “wilcoxauc” function from the presto package (version 1.0.0) or “FindMarkers” function in Seurat to compare groups based on the specified metadata (e.g., cluster, time point). Genes were ranked by the AUC statistic or log_2_ fold change in descending order, and these rankings were used as input for the enrichment analysis. Enrichment scores and normalized enrichment scores (NES) were computed for each pathway, and significance was assessed using adjusted *P*-values (Benjamini-Hochberg correction). Pathways with adjusted *P*-values < 0.05 were considered significantly enriched.

### Analysis of signaling pathway activity

To infer signaling pathway activity from transcriptomic data, we used the decoupleR package (version 2.7.1)(*67*), which implements a knowledge-based model of Pathway RespOnsive GENes for activity inference (PROGENy)(*68*) derived from a large compendium of perturbation experiments. Signaling pathway activity scores at the single-cell level were computed by employing the linear model. The resulting pathway activity scores were averaged within each cluster, and these cluster-level mean scores were used for visualization.

### Inferring the cellular origins of CCR7^+^ DCs and ISG⁺ DCs

To investigate the cellular origins of CCR7^+^ DCs and ISG⁺ DCs, we applied the SingleR algorithm(*33*). This tool infers cell identity by calculating Spearman correlation coefficients between the transcriptomes of individual cells and those within a reference dataset. In this analysis, we used transcriptomic profiles of CCR7⁺ISG15^hi^CD1C⁺ and CCR7⁺ISG15^hi^CADM1⁺ subsets as reference to assign potential origins of mature DCs across different tumor types. Additionally, cDC2 subsets were employed as references to explore lineage relationships between ISG⁺ DCs and cDC2-derived CCR7^+^ DCs.

### Generating the reference scFlex-seq dataset

The single-cell reference atlas(*35*) included level 1 and level 2 cell type annotations, which were previously manually classified (**Supplemental Figure 6A**). Level 1 represented broad cell types, while level 2 denoted more detailed cell types generated through repeated clustering within each level 1 cluster. We performed a clustering process on the level 1 myeloid cluster, and identified clusters positive for *FLT3*, *ZBTB46*, and *ITGAX*, and annotated as cDC1, cDC2, ISG⁺ DCs, and CCR7^+^ DCs based on DEGs. To investigate the phenotypes of T cells co-localizing with DCs, cells within the CD4 T cell and CD8 T cell clusters in Level2 annotation were mapped onto the pre-defined human CD4 TIL atlas and human CD8 TIL atlas, respectively, which is generated by projecTILs framework (version 3.3.1)(*69*) (**Supplemental Figure 6**). Since ProjecTILs defines CD8.EX based upon the expression level of five markers (*TOX, PDCD1, LAG3, TIGIT*, and *HAVCR2*), CD8.EX potentially encompass heterogeneous populations, such as activated, effector-like exhausted, and dysfunctional T cells. These additional labels for DCs and T cells were integrated into the level 2 annotations. We then used this annotated single-cell dataset as a reference to deconvolve the Visium HD data, assigning a homogeneous set of cell type labels across the different samples.

### Cell type deconvolution of Visium HD spatial transcriptomics data

To assign cell types in Visium HD spatial transcriptomics data using the scFlex-seq reference derived from serial sections (as detailed above), we applied the Robust Cell Type Decomposition (RCTD) algorithm(*70*), implemented in the spacexr R package (version 2.2.1). RCTD allows for spatial pixels to be assigned sparse mixtures of the reference cell types. Following conversion of the Visium HD spatial data into a Seurat object, we utilized the sketch-based analysis feature(*71*) implemented in Seurat v5, which strategically subsamples approximately 50,000 representative cells while preserving the diversity of both abundant and rare cell types. This analytical step was performed to optimize computational resources while ensuring a comprehensive representation of cellular diversity within the dataset. The mapping was performed using RCTD in doublet-aware mode, based on 8 µm spatial bins. After running RCTD, the predicted cell type classifications were projected from the sketch to the full spatial dataset using the “ProjectData” function. The deconvolved Visium HD data provided a high-resolution spatial map of cell types consistent with tissue morphology and known gene expression markers such as *PIGR* (goblet cells and enterocytes), *CEACAM6* (tumor cells), and *COL1A1* (fibroblasts) (**Supplemental Figures 7A-C**).

### Cell segmentation and bin-to-cell conversion of Visium HD

To perform the distance-based analysis, we reconstructed bins into cells by leveraging the recently published pipeline Bin2cell(*72*). Cell segmentation was performed using the bin2cell workflow in a Python environment (version 3.12.4), which reconstructs cells by integrating high-resolution (2 μm bin) gene expression data and morphology-based image segmentation. Briefly, 2 μm bins were assigned to individual cells based on nuclei segmentation obtained from H&E-stained images using StarDist (version 0.9.1), a deep learning-based tool for nuclei detection. To refine segmentation, identified nuclei were expanded to neighboring unlabeled bins, enabling more comprehensive cell coverage. In parallel, gene expression profiles were used for secondary cell assignment, allowing for the inclusion of cells in regions lacking detectable nuclei. Reconstructed cells were filtered based on the following criteria: number of bins per cell > 5 and < 100. For cell type identification, unsupervised clustering was performed using the Seurat pipeline, incorporating an additional quality control step. Clusters were annotated using results from spot deconvolution as reference.

### Analysis of putative spatial cell-cell interactions

To investigate spatially proximal ligand-receptor interactions, we utilized CellChat v2 (version 2.1.2) (*73*). The sketch-based Visium HD spatial transcriptomics data (8 µm bins) were imported into CellChat, and the CellChatDB was used as the reference ligand-receptor interaction database. All preprocessing steps, including normalization and identification of overexpressed genes, were performed separately for each sample. Communication probabilities between cell types were computed using the “computeCommunProb” function with the n_contact.range parameter set to 30, which restricts cell-cell contact dependent interaction inference to spatially neighboring cells. To reduce noise potentially introduced by spatial bins containing mixed transcriptomes from multiple cells, we excluded from visualization any inferred interactions involving cell types with zero expression of either the ligand or receptor gene in the scFlex-seq reference dataset.

### Statistical analysis

The two-sided Wilcoxon rank sum test was used for comparisons between two groups. Differences between multiple variables were assessed using a one-way analysis of variance (ANOVA) with Tukey’s multiple comparisons test. For scRNA-seq analysis, adjusted *P*-values for differential expression gene analysis were calculated using a two-sided non-parametric Wilcoxon rank sum test with Bonferroni correction. Statistical analysis was performed using R software (version 4.4.0). *P* < 0.05 was considered a significant difference.

## Acknowledgments

We thank Dr. Mauro Di Pilato (University of Texas MD Anderson Cancer Center) for insightful discussions and critical feedback on the manuscript. We thank Dr. Ruan FV Medrano and Dr. Robert D. Schreiber (Washington University) for their assistance with the design of the CODEX marker panel. We thank Dr. Patricia Castro (Baylor College of Medicine) for her assistance with preparation of CODEX fresh frozen tissue sections. We also thank Ms. Amber Kao (University of Texas MD Anderson Cancer Center) for her helpful suggestions regarding CODEX-related analyses. We acknowledge the support of the High-Performance Computing for Research facility at the University of Texas MD Anderson Cancer Center for providing computational resources. Graphics were created with BioRender.com.

## Author contributions

Conceptualization, T.M., S.S.W., and M.M.G.; data curation, T.M., and S.K.,; investigation, T.M., S.K., A.S.S., A.S., J.E.P., and M.M.G.; CODEX multiplexed imaging, S.K.; visualization, T.M. S.K., S.H., K.H.H., and K.C.; methodology, T.M., S.K., S.H., Y.Y.Y., B.Z., V.M., S.J., L.W., G.P.N., S.S.W., and M.M.G.; data analysis, T.M., S.K., S.H., Y.Y.Y., B.Z., V.M., S.J., K.H.H., K.C., L.W. and G.P.N.; writing – original draft, T.M., S.K., S.S.W., and M.M.G.; writing – review & editing, S.K., A.S.S., A.S., J.E.P., K.H.H., K.C., S.S.W., and M.M.G.; supervision, S.S.W., and M.M.G.; project administration, S.S.W., and M.M.G.

## Data and materials availability

All data associated with this study are present in the paper or the Supplemental Materials. The processed integrated human or mouse scRNA-seq object of tumor-infiltrating DCs and processed CODEX AnnData files are publicly available on https://zenodo.org with the DOI: 10.5281/zenodo.18157069. Codes used for data analysis are available at GitHub (https://github.com/T-Minowa/TIDCmap). All other data and materials generated in this study are available from the corresponding authors upon reasonable request.

---

**Supplemental Fig. 1.**
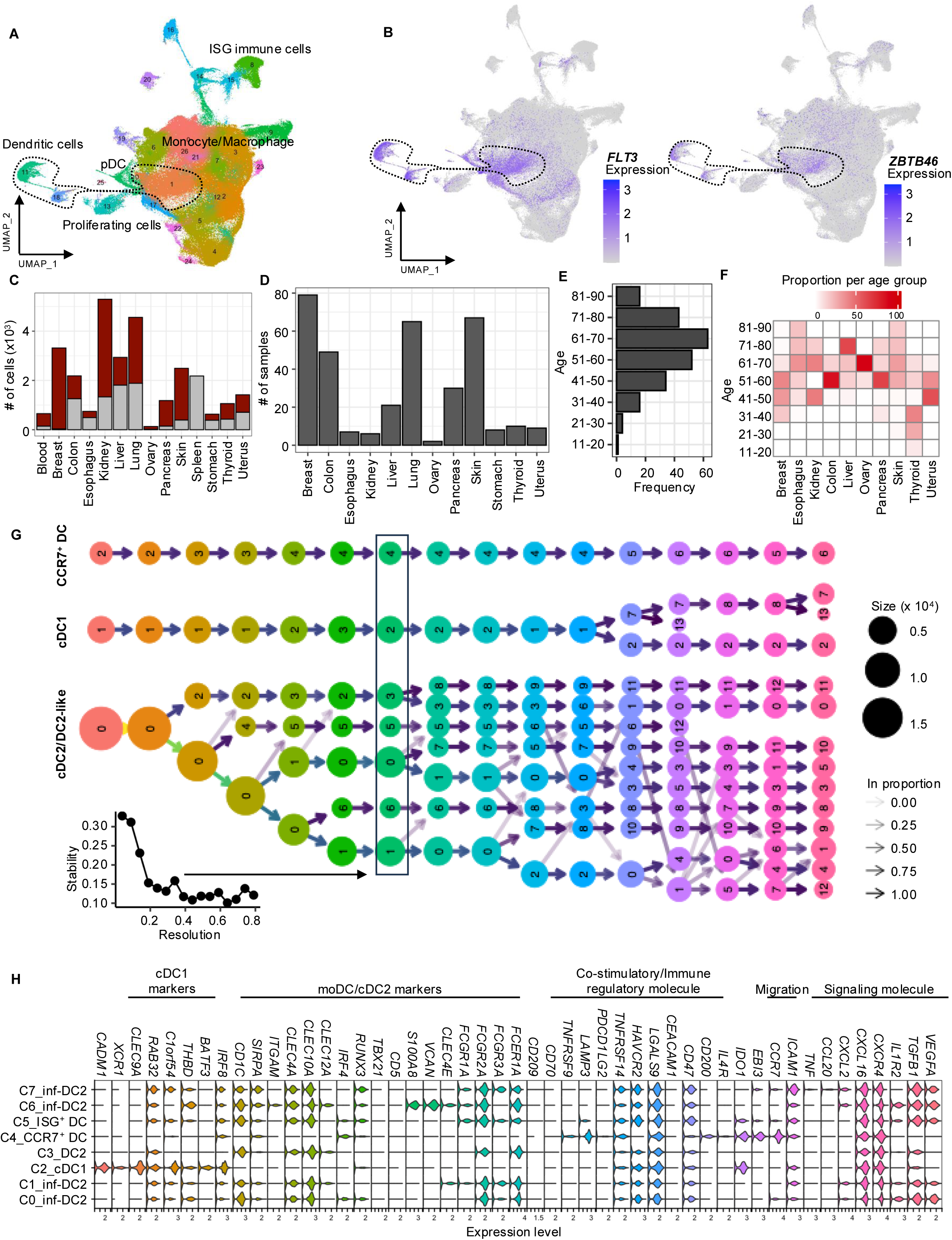
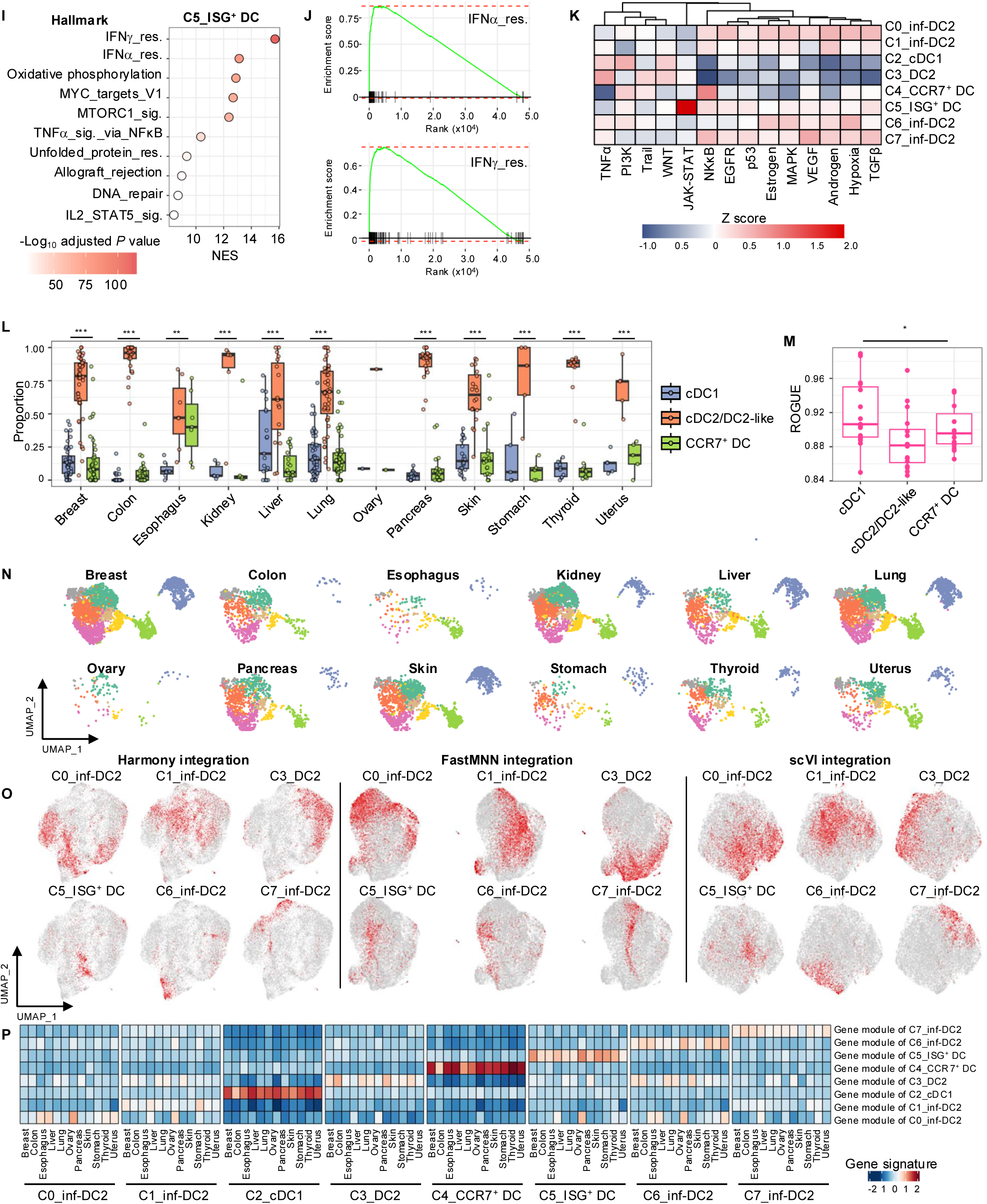
Construction of tumor-infiltrating DC atlas and characterization of cDC subsets in human. **(A)** Integrated UMAP visualization of monocytes, macrophages, DCs, and other immune cells derived from tumor, adjacent normal tissue, spleen, and blood. Clusters are denoted by colors. **(B)** UMAP plots showing the expression of *FLT3* and *ZBTB46*, lineage markers defining conventional DCs, with color intensity indicating expression level. **(C)** Bar plot showing the number of DCs identified in each tissue. Red bars represent tumor-infiltrating DCs, and grey bars represent DCs from the normal tissue **(D)** Bar plot showing the number of samples analyzed for each tissue type. **(E)** Histogram representing the age distribution of patients included in the tumor-infiltrating DCs. **(F)** Heatmap showing the proportion of samples from each age group for each tissue type. **(G)** Clustree plot showing clustering relationships across resolutions (k = 0.05-0.8). The clustering resolution selected for downstream analyses is highlighted by the enclosed square and indicated by an arrow from the bar plot displaying mean SC3 stability across resolutions. **(H)** Violin plot showing the expression of marker genes for tumor-infiltrating DCs across clusters. **(I)** Dot plot illustrating the top 10 highest normalized enrichment scores (NES) of MSigDB hallmark pathways in C5_ISG⁺ DC compared to other DC clusters. The dot color indicates the adjusted *P*-value. **(J)** GSEA plots for the Hallmark IFN-a (top) and IFN-g (bottom) response pathways in C5_ISG⁺ DC. **(K)** Heatmap of the normalized signaling pathway activity for tumor-infiltrating DCs. **(L)** Boxplot showing the tissue-specific proportion of tumor-infiltrating DC subtypes. Each dot represents the individual fractions for samples. Samples with fewer than 10 cells were excluded. The lines within the box and whiskers indicate the median and quartiles, respectively. *P*-values (\**P* < 0.05, \*\**P* < 0.01, \*\*\**P* < 0.001, \*\*\*\**P* < 0.0001) were calculated using one-way ANOVA. **(M)** Boxplot showing the score of ROGUE for tumor-infiltrating DC subtypes. The lines within the box and whiskers indicate the median and quartiles, respectively. *P*-values (\**P* < 0.05) were calculated using one-way ANOVA. **(N)** UMAP plots for each tissue type, showing the distribution of tumor-infiltrating DCs subsets across tissues. Clusters are denoted by colors. **(O)** UMAP plots of cDC2 population generated from Harmony (left), FastMNN (middle), and scVI (right) integration. Red dots indicate cells corresponding to each cluster defined by the reciprocal PCA integration method. **(P)** Heatmaps showing signature scores for tumor-infiltrating DC clusters across cancer types. Normalized average module scores for each cluster and cancer type are displayed.

**Supplemental Fig. 2.**
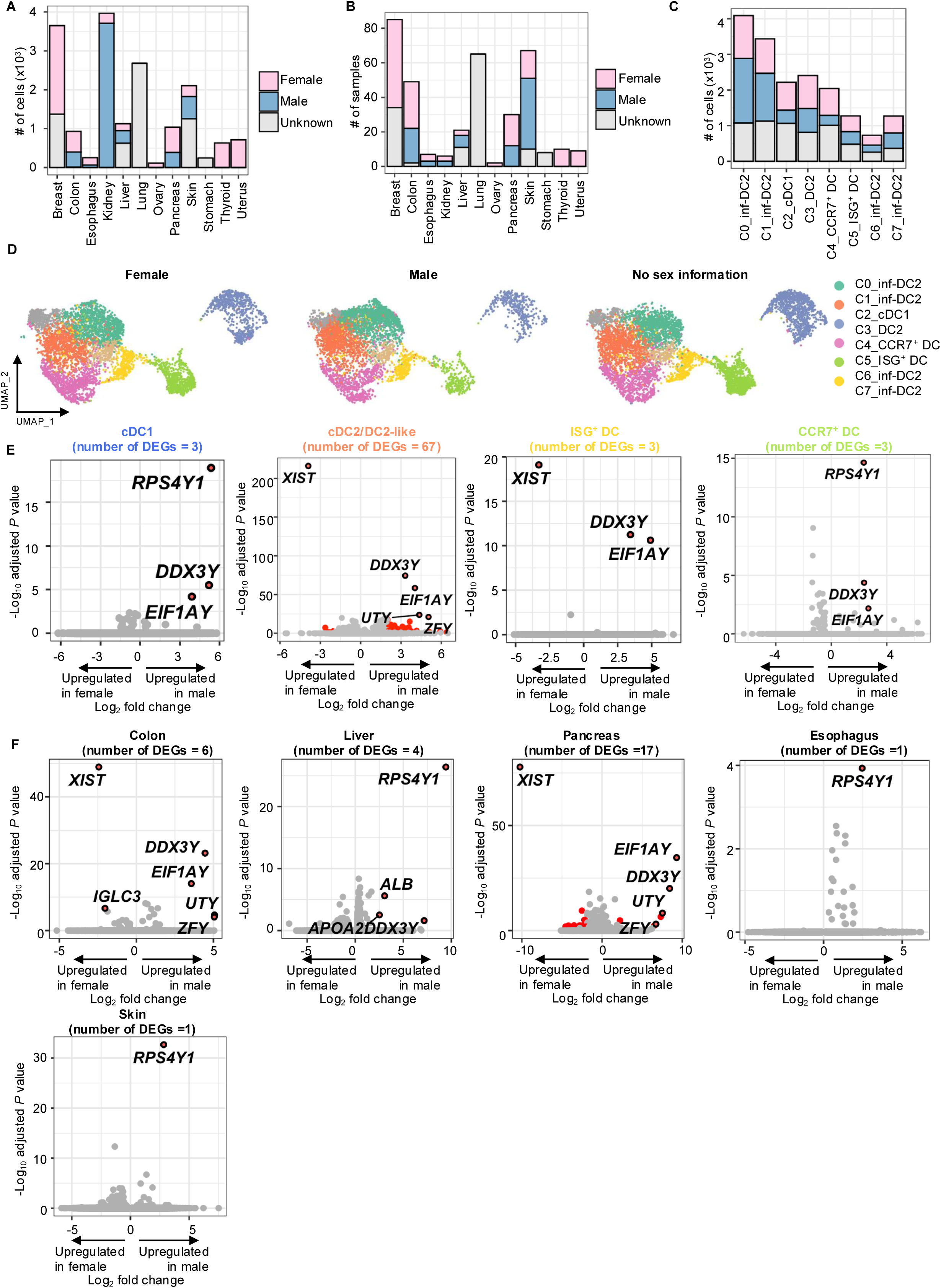
Sex-based analysis of tumor-infiltrating DCs across tissues. **(A)** Bar plot showing the number of tumor-infiltrating DCs for each tissue type, stratified by sex. **(B)** Bar plot showing the number of samples for each tissue type, stratified by sex. **(C)** Bar plot showing the number of tumor-infiltrating DCs in each cluster, stratified by sex. **(D)** UMAP plots illustrating the distribution of tumor-infiltrating DCs based on sex. Clusters are denoted by colors. **(E)** Volcano plots showing sex-associated DEGs within individual DC subtypes. Significantly differentially expressed transcripts (log_2_FC > 2, adjusted *P* value < 0.05) are shown in red. Statistical analysis was performed using a two-sided nonparametric Wilcoxon rank sum test with Bonferroni’s correction. **(F)** Volcano plots showing sex-associated DEGs of cDC2 cluster within individual tissue types. Significantly differentially expressed transcripts (log_2_FC > 2, adjusted *P* value < 0.05) are shown in red. Statistical analysis was performed using a two-sided nonparametric Wilcoxon rank sum test with Bonferroni’s correction.

**Supplemental Fig. 3.**
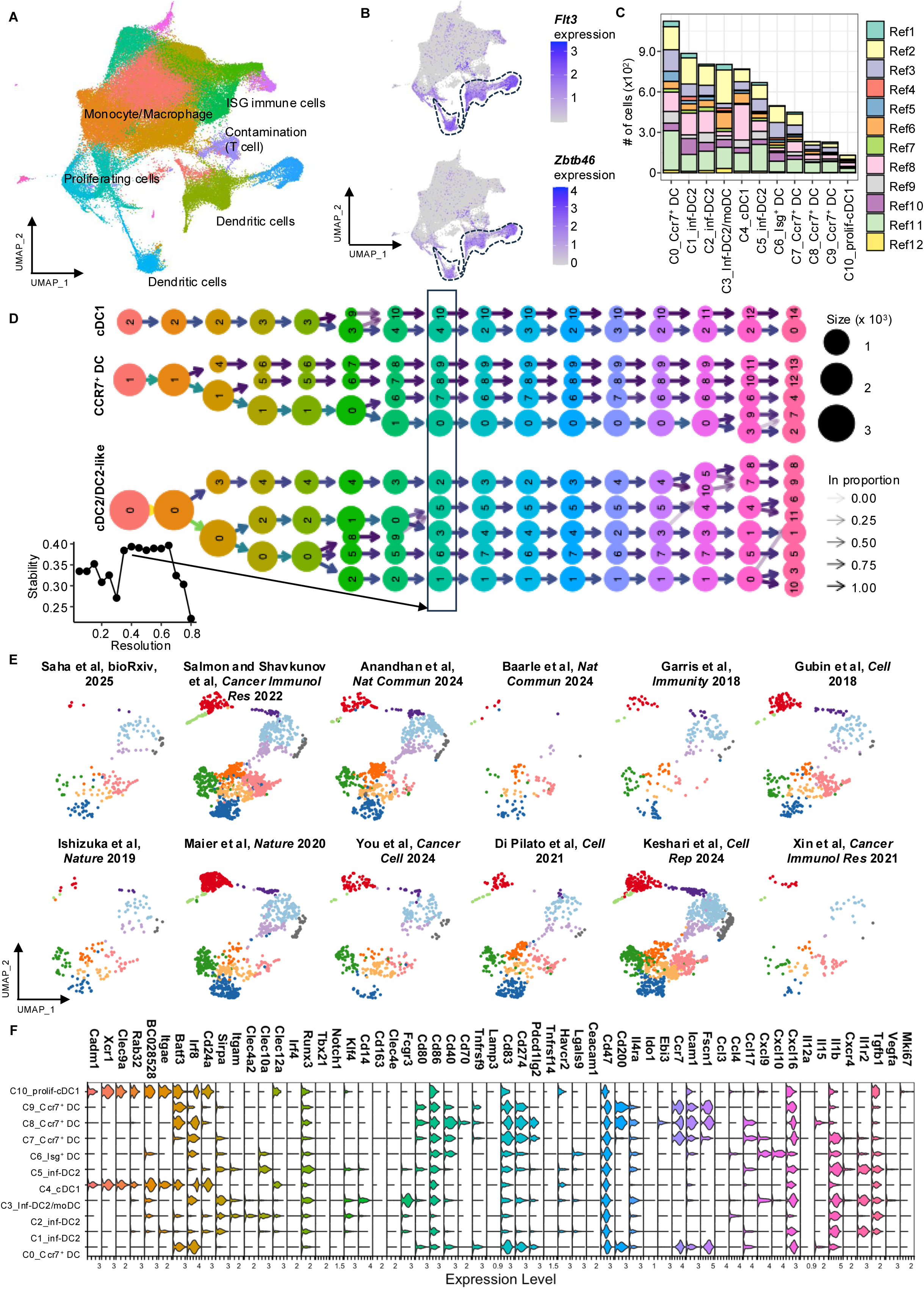
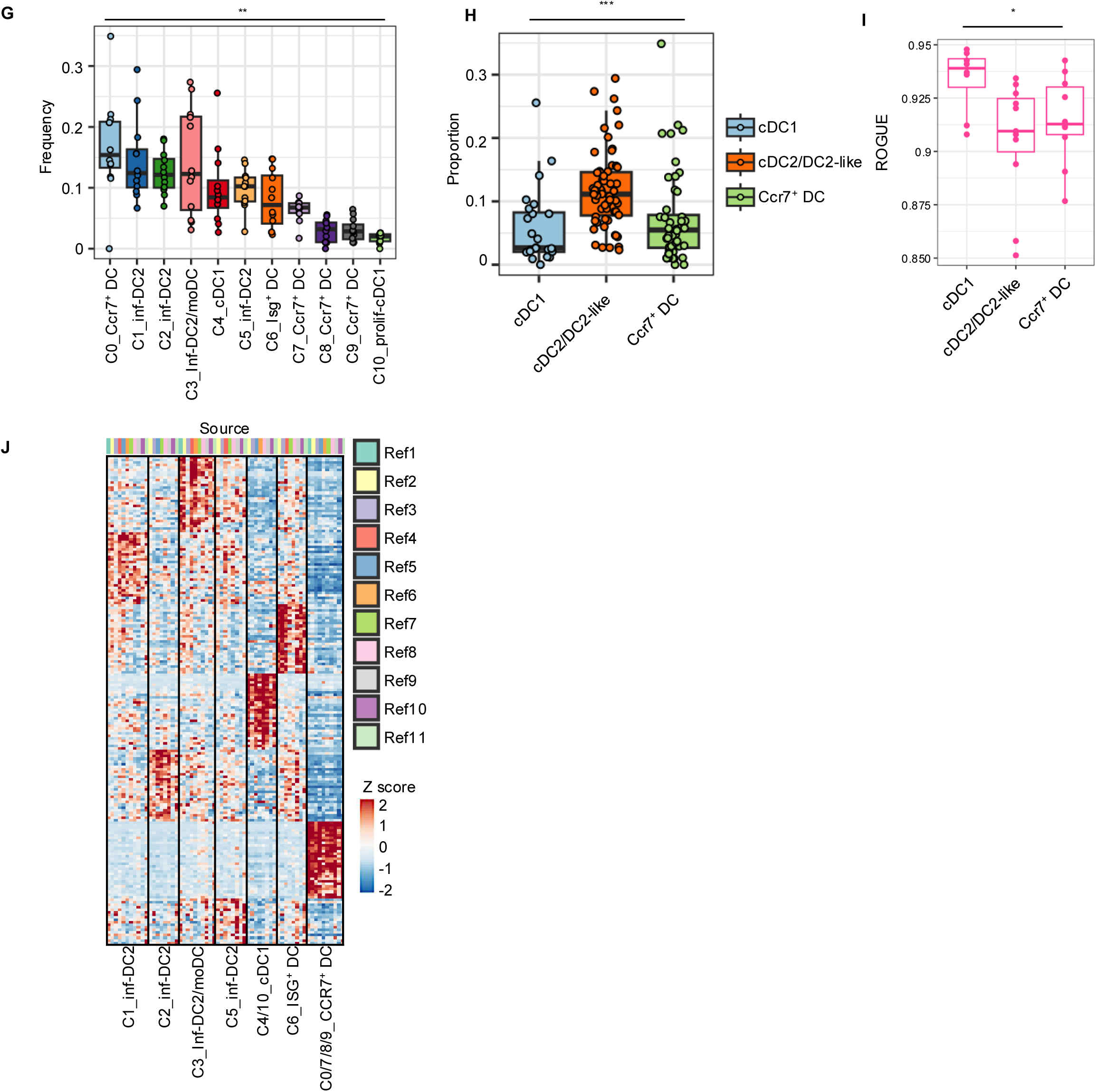
Construction and characterization of a tumor-infiltrating DC atlas in mouse. **(A)** Integrated UMAP visualization of monocytes, macrophages, DCs, and other immune cells derived from murine tumors. Clusters are denoted by colors. **(B)** UMAP plots showing the expression of lineage markers defining conventional DCs, with color intensity indicating expression level. **(C)** Bar plot showing the number of tumor-infiltrating DCs in each cluster, with colors indicating the contribution from each cohort. **(D)** Clustree plot showing clustering relationships across resolutions (k = 0.05-0.8). The clustering resolution selected for downstream analyses is highlighted by the enclosed square and indicated by an arrow from the bar plot displaying mean SC3 stability across resolutions. **(E)** UMAP plots showing the distribution of tumor-infiltrating DCs from each individual cohort within the integrated mouse DC atlas. **(F)** Violin plot showing the expression of selected marker genes for mouse tumor-infiltrating DC clusters. **(G) (H)** Boxplot showing the frequency of each DC cluster across different cohorts. The lines within the box and whiskers indicate the median and quartiles, respectively. *P*-values (\*\**P* < 0.01) were calculated using one-way ANOVA. **(H)** Boxplot showing the ROGUE score, an entropy-based metric for cell cluster purity, for major DC subtypes. The lines within the box and whiskers indicate the median and quartiles, respectively. *P*-values (\*\**P* < 0.01) were calculated using one-way ANOVA. **(I)** Heatmap showing normalized gene expression in mouse tumor-infiltrating DC clusters across cohort. The heatmap displays the top 30 DEGs with the highest log_2_ fold change and a detection percentage (pct.1) above 0.5, as identified by DEG analysis.

**Supplemental Fig. 4.**
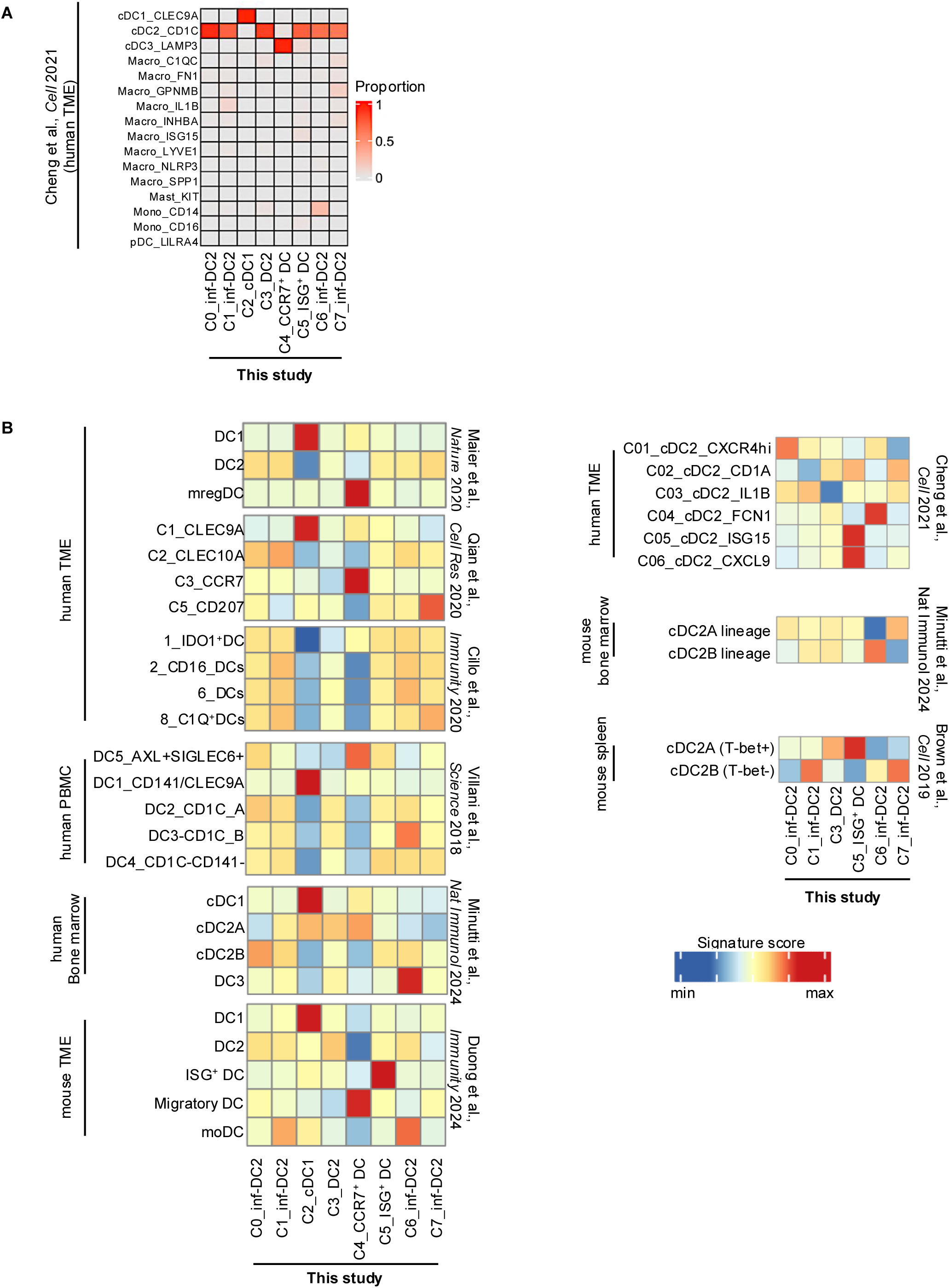
Inter-cohort comparison of DC clusters. **(A)** Heatmap representing a confusion matrix between independent cell type annotations (this study and Cheng et al.(*14*)). **(B)** Heatmap depicting the signature scores in tumor-infiltrating DCs of this study (rows) using reference gene signatures (columns) derived from the analyses by from Qian *et al*.(*19*), Cillo *et al.*(*75*), Villani *et al*.(*27*), Maier *et al.*(*12*), Minutti *et al.*(*76*), Brown *et al.*(*77*), and Cheng *et al.*(*14*). Signature scores calculated in individual cells were averaged for each cluster and were used to visualize the results.

**Supplemental Fig. 5.**
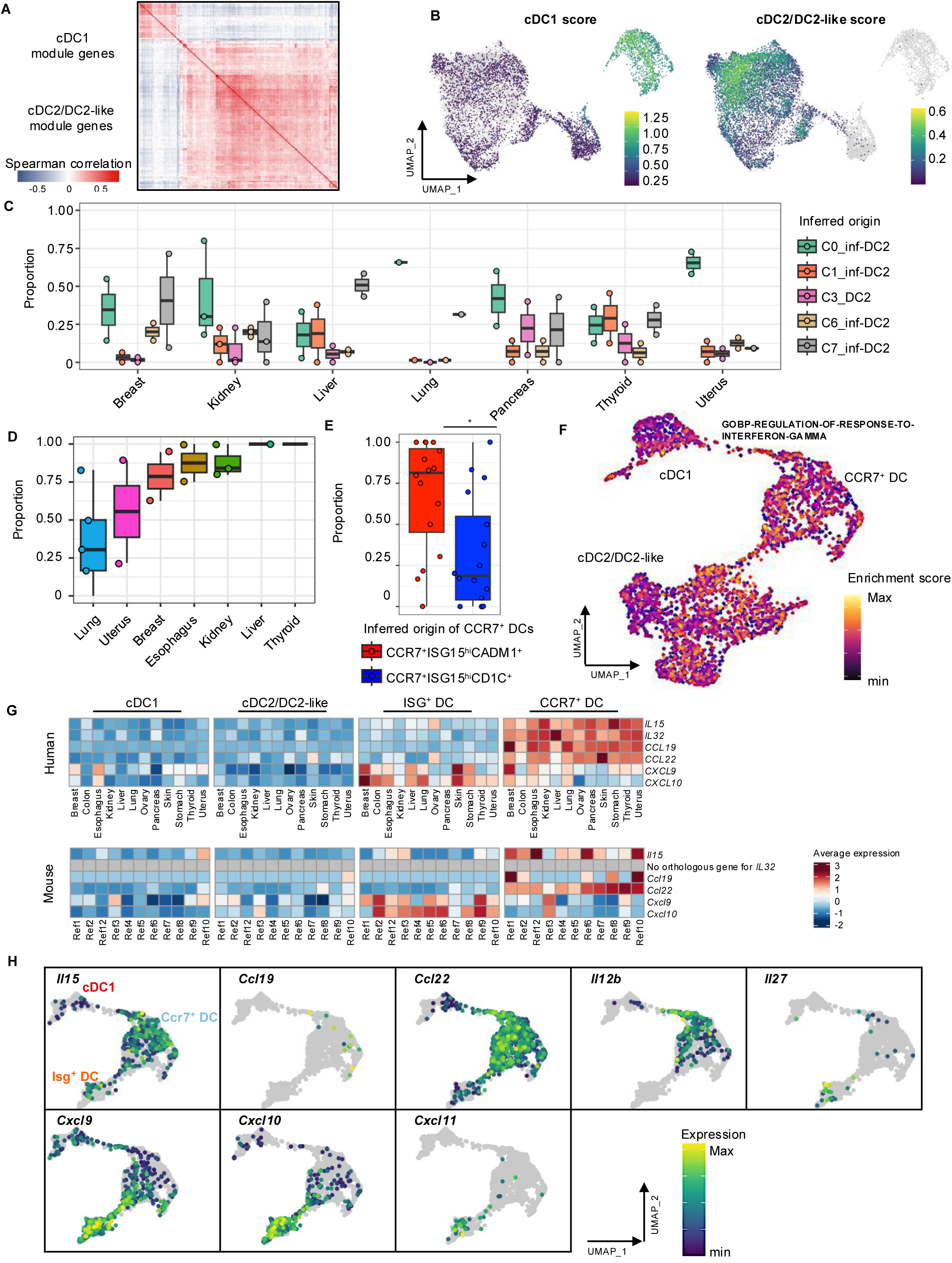
Functional characterization and lineage analysis of human ISG⁺ DCs and CCR7^+^ DCs. **(A)** Heatmap showing the Spearman correlation of module genes between cDC1 and cDC2 clusters. **(B)** UMAP plots showing cDC1 and cDC2 signature scores projected onto the integrated human DC atlas. **(C)** Boxplot showing the inferred origin fractions of C5_ISG⁺ DC from different cDC2 subtypes across cancer types. Samples with more than three cells in the C5_ISG⁺ DC cluster were included; samples lacking cDC2 subclusters were excluded. The lines within the box and whiskers indicate the median and quartiles, respectively. Each dot indicated samples. **(D)** Boxplot showing the inferred fractions of CCR7^+^ISG15^hi^CADM1^+^-derived mature DCs across cancer types. Samples with more than two cells in the C4_CCR7^+^ DC cluster were included; samples lacking CCR7⁺ISG15^hi^CADM1⁺ or CCR7⁺ISG15^hi^CD1C⁺ subclusters were excluded. The lines within the box and whiskers indicate the median and quartiles, respectively. Each dot indicated samples. **(E)** Boxplot showing the proportion of inferred cDC1-derived (CCR7^+^ISG15^hi^CADM1^+^) and cDC2-derived (CCR7^+^ISG15^hi^CD1C^+^) cells within the C4_CCR7^+^ DC cluster. The lines within the box and whiskers indicate the median and quartiles, respectively. Each dot indicated samples. *P*-values (\**P* < 0.05) were calculated using the Wilcoxon rank-sum test. **(F)** UMAP displaying *Isg15* expression in C6_ISG⁺ DC, C4_cDC1, and CCR7^+^ DC from mouse dataset. **(G)** Heatmaps showing normalized expression of genes encoding cytokines and chemokines in human DCs (top) and mouse DCs (bottom) across cancer types in humans and across studies in mice. **(H)** UMAP displaying normalized gene expression in C6_ISG⁺ DC, C4_cDC1, and CCR7^+^ DC clusters from mouse dataset.

**Supplemental Fig. 6.**
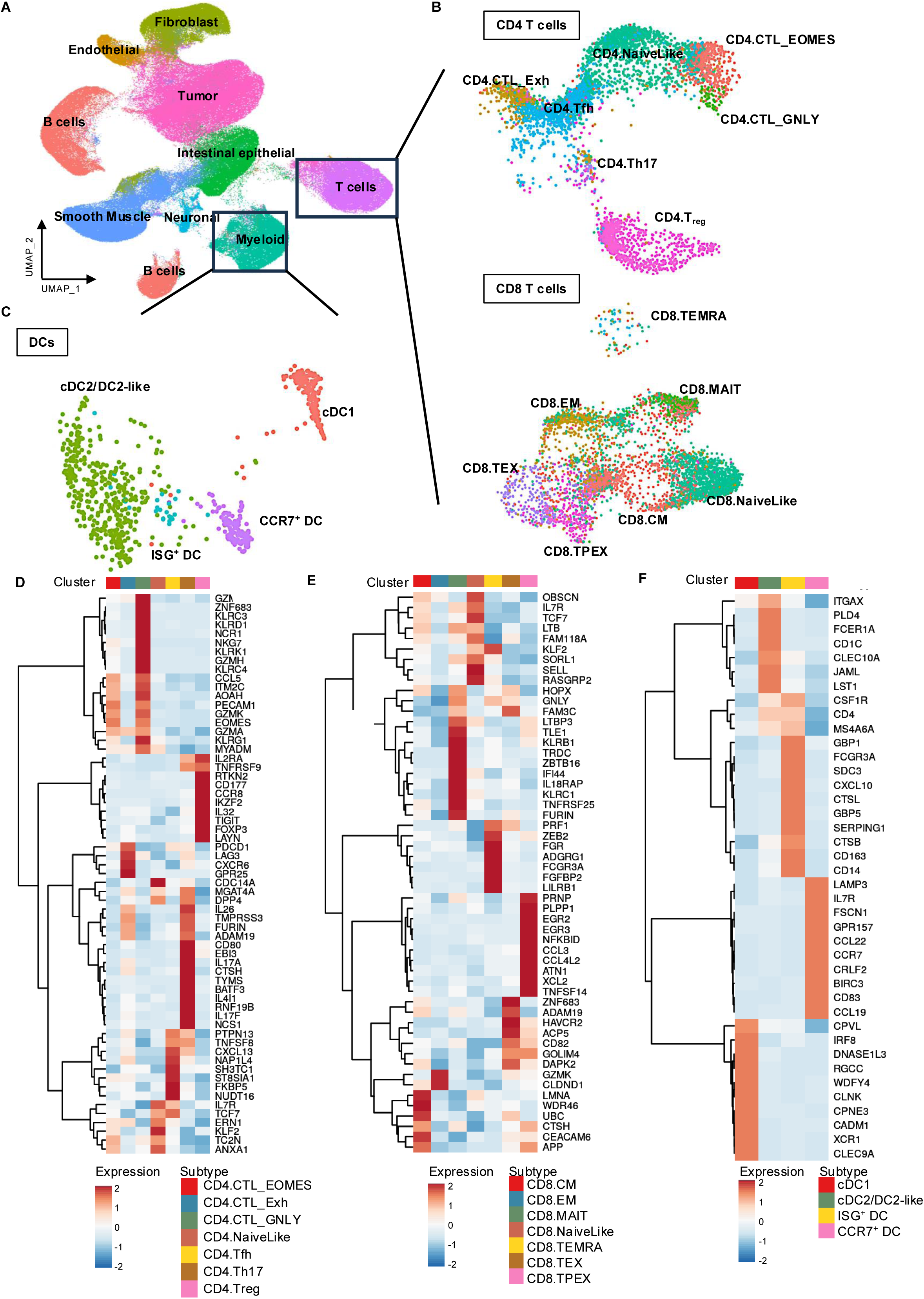
Reference single-cell dataset for spatial transcriptomics deconvolution. **(A)** UMAP visualization of all cell types in the scFlex-seq reference dataset from human colorectal cancer (CRC). Level 1 cell type annotations are labeled. **(B)** Mapping of T cells to CD4 and CD8 TIL references using projecTILs framework(*69*). **(C)** UMAP plot showing DCs after subclustering for annotating cDC1, cDC2, ISG⁺ DC, and CCR7^+^ DC. **(D) (E) (F)** Heatmaps showing normalized expression of marker genes for the annotated cell subtypes. Marker genes were selected as the top 10 upregulated DEGs with the highest log_2_ fold change in each cluster.

**Supplemental Fig. 7.**
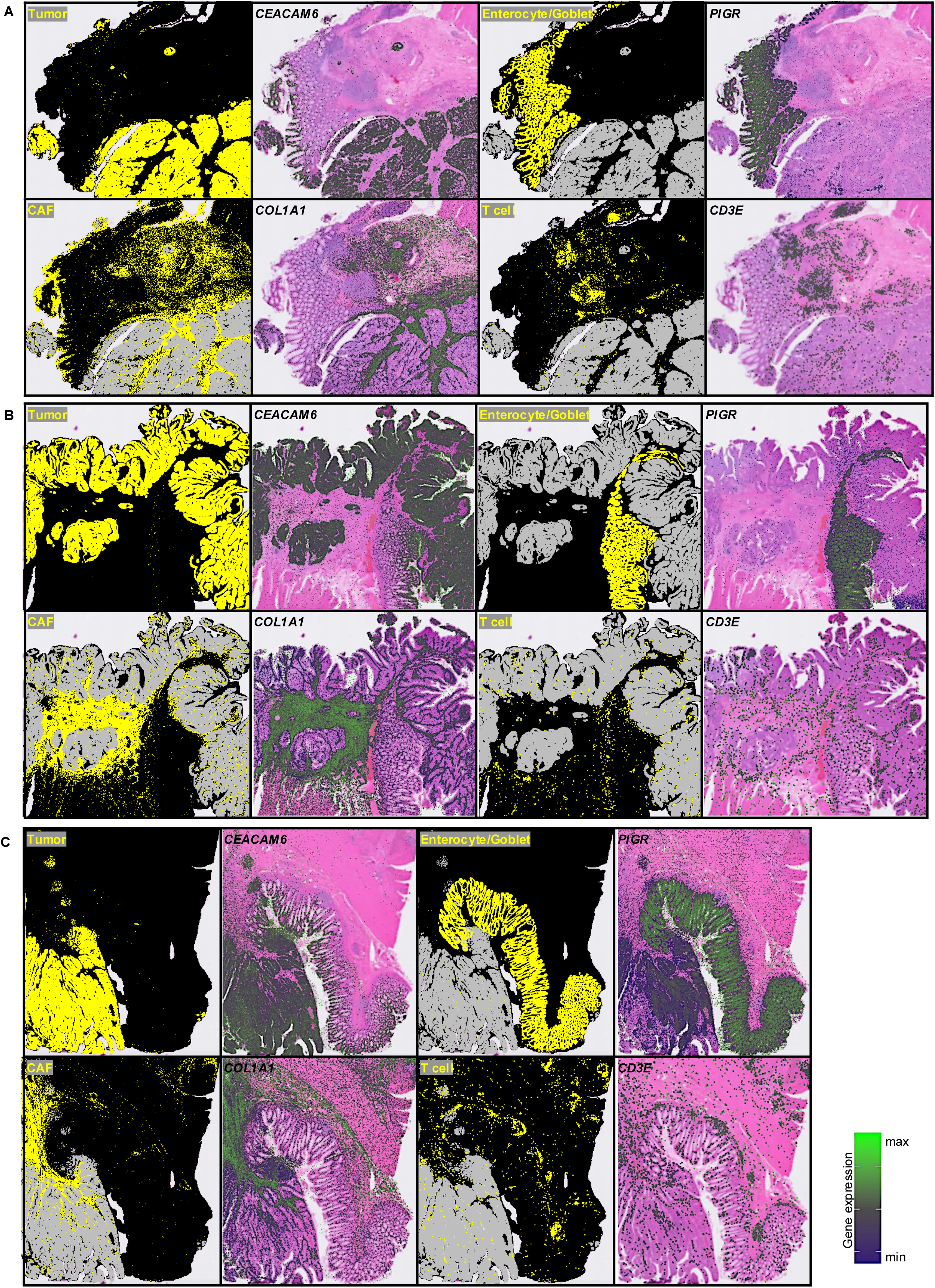
Validation of cell type deconvolution in spatial transcriptomics data. **(A) (B) (C)** Spatial plots from three human colorectal cancer (CRC) samples (CRC patient1, CRC patient2, and CRC patient5) comparing the distribution of deconvoluted cell types (Tumor, CAF, Enterocyte/Goblet, T cell) with the spatial expression of their canonical marker genes (*CEACAM6*, *COL1A1*, *PIGR*, and *CD3E*, respectively).

**Supplemental Fig. 8.**
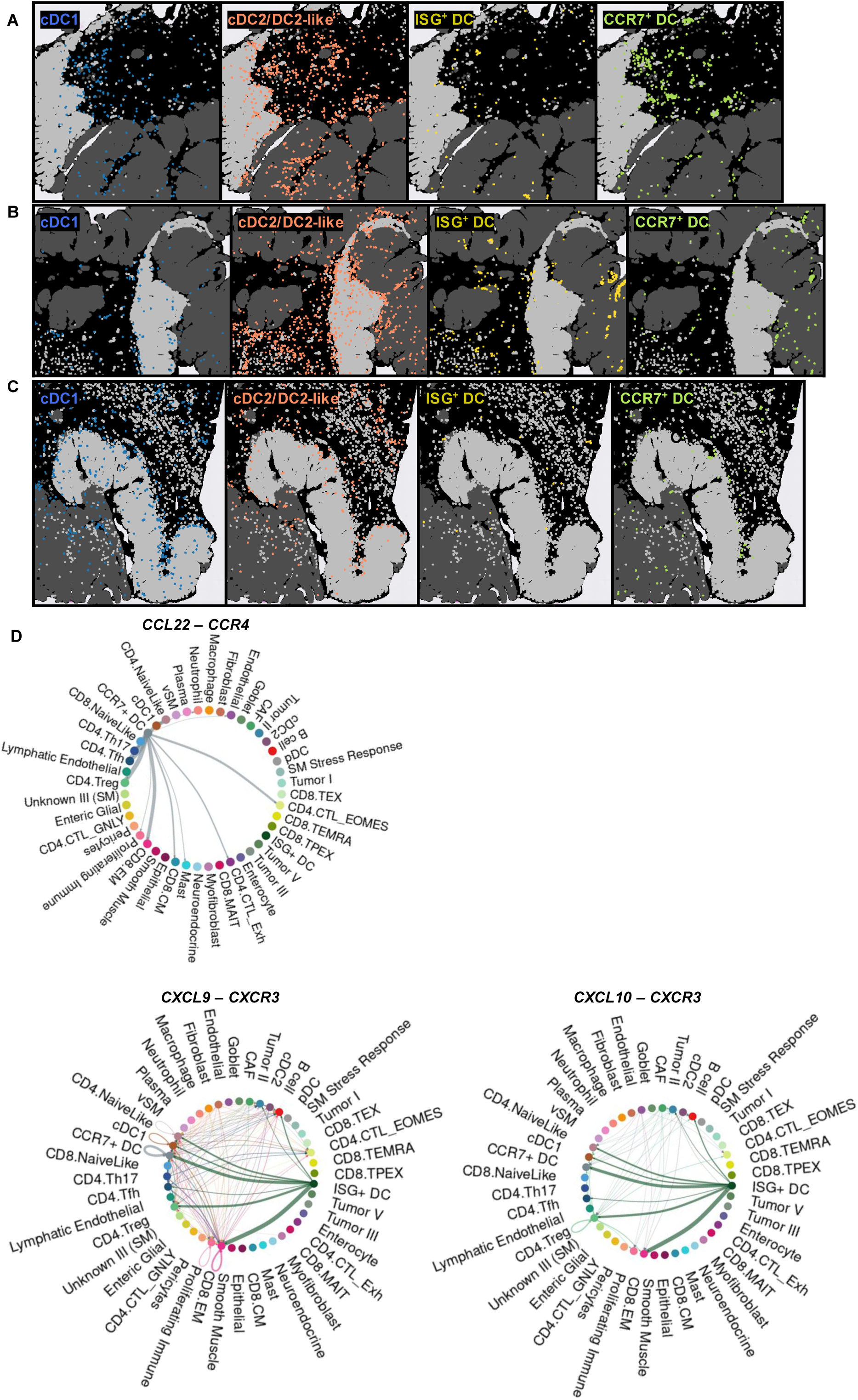
Spatial distribution and proximity analysis of DC. **(A) (B) (C)** Spatial mapping of tumor-infiltrating DCs from three human colorectal cancer (CRC) samples of (A) CRC patient1, (B) CRCP2, and (C) CRCP5. **(D)** Chord diagram visualizing cell-cell communication via the CCL22-CCR4 and CXCL9/10-CXCR3 ligand-receptor axes. For the CCL22-CCR4 axis, only CCL22 signals originating from CCR7⁺ DCs are shown, whereas for the CXCL9/10-CXCR3 axis, ligand signals from all validated source cell types are visualized. Cells are included as ligand sources or targets only when expression of the corresponding ligands or receptors is supported by the scFlex-seq reference dataset.

**Supplemental Fig. 9.**
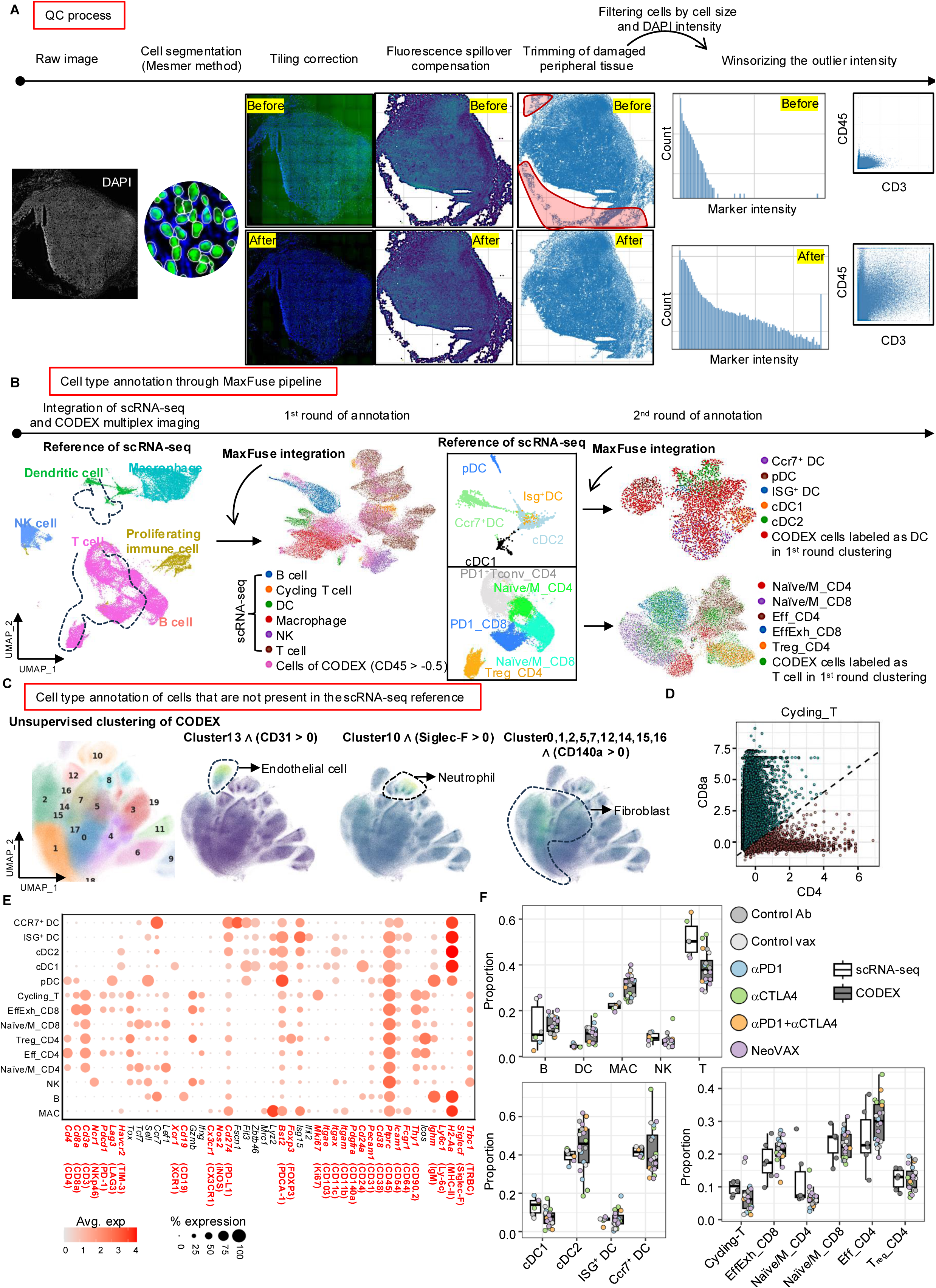
Quality control, integration, and cell-type annotation of multiplexed CODEX data. **(A)** Overview of the quality control (QC) workflow applied to raw CODEX images, with representative examples shown before and after each QC step. **(B)** Cell-type annotation of CODEX multiplexed imaging data through integration with scRNA-seq reference datasets using the MaxFuse algorithm. **(C)** Manual annotation of endothelial cells, neutrophils, and fibroblasts based on unsupervised clustering of CODEX cells and marker expression. **(D)** Scatter plot showing normalized protein intensities of CD4 and CD8a in cells annotated as cycling T cells. The diagonal line subdivides cycling T cells into CD4 cycling T cells and CD8 cycling T cells. **(E)** Dot plot showing gene expression patterns in the reference scRNA-seq dataset. The first and second rows of the column labels indicate gene names and corresponding protein names, respectively. Gene names that correspond to protein markers used in CODEX are highlighted in red. **(F)** Box plots showing cell-type proportions derived from scRNA-seq and CODEX datasets across treatment conditions. The lines within the box and whiskers indicate the median and quartiles, respectively.

**Supplemental Fig. 10.**
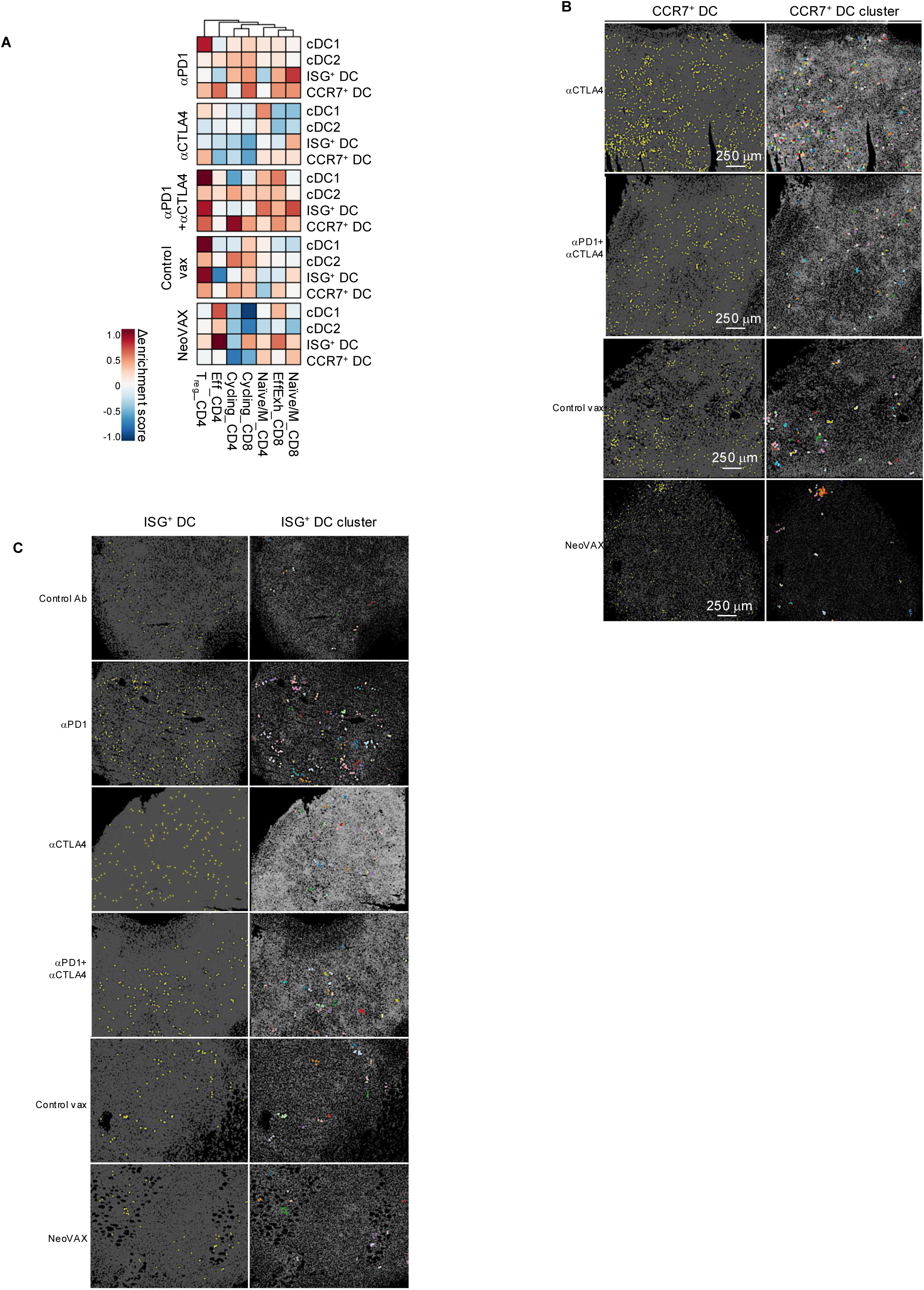
Therapy-dependent modulation of DC-T cell neighborhood enrichment and DC clustering patterns. **(A)** Heatmaps showing changes in neighborhood enrichment scores between treatment conditions and the corresponding control groups for the spatial proximity of DC subtypes and T cell phenotypes. Control antibody treated samples were used as the baseline for ICT-treated groups, and control vax samples were used as the baseline for the NeoVAX group. **(B)** Representative spatial maps illustrating the distribution of CCR7⁺ DCs (left) and CCR7⁺ DC clusters (right) across treatment conditions. Representative spatial maps illustrating the distribution of ISG⁺ DCs (left) and ISG⁺ DC clusters (right) across treatment conditions.

**Supplemental Fig. 11.**
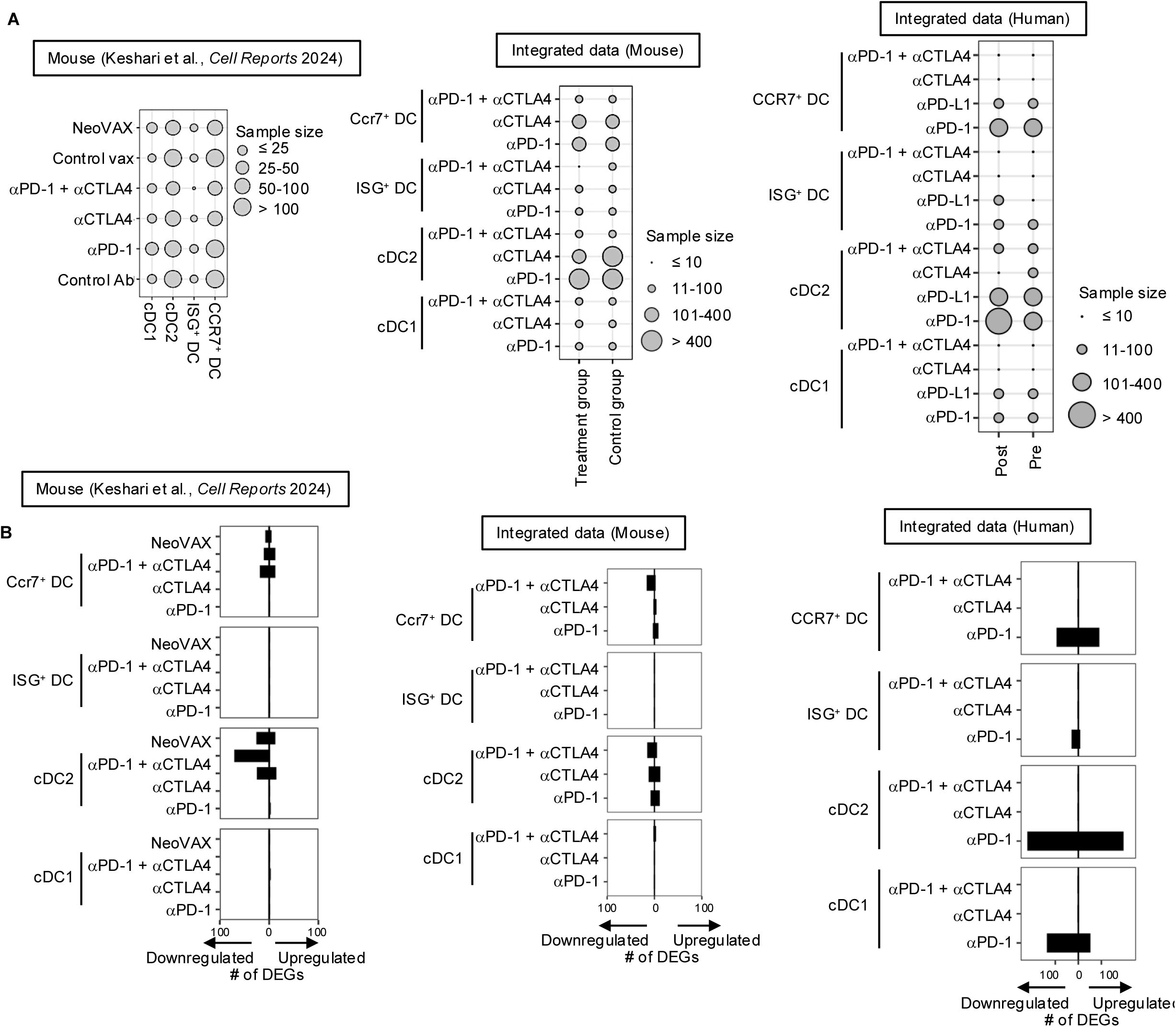
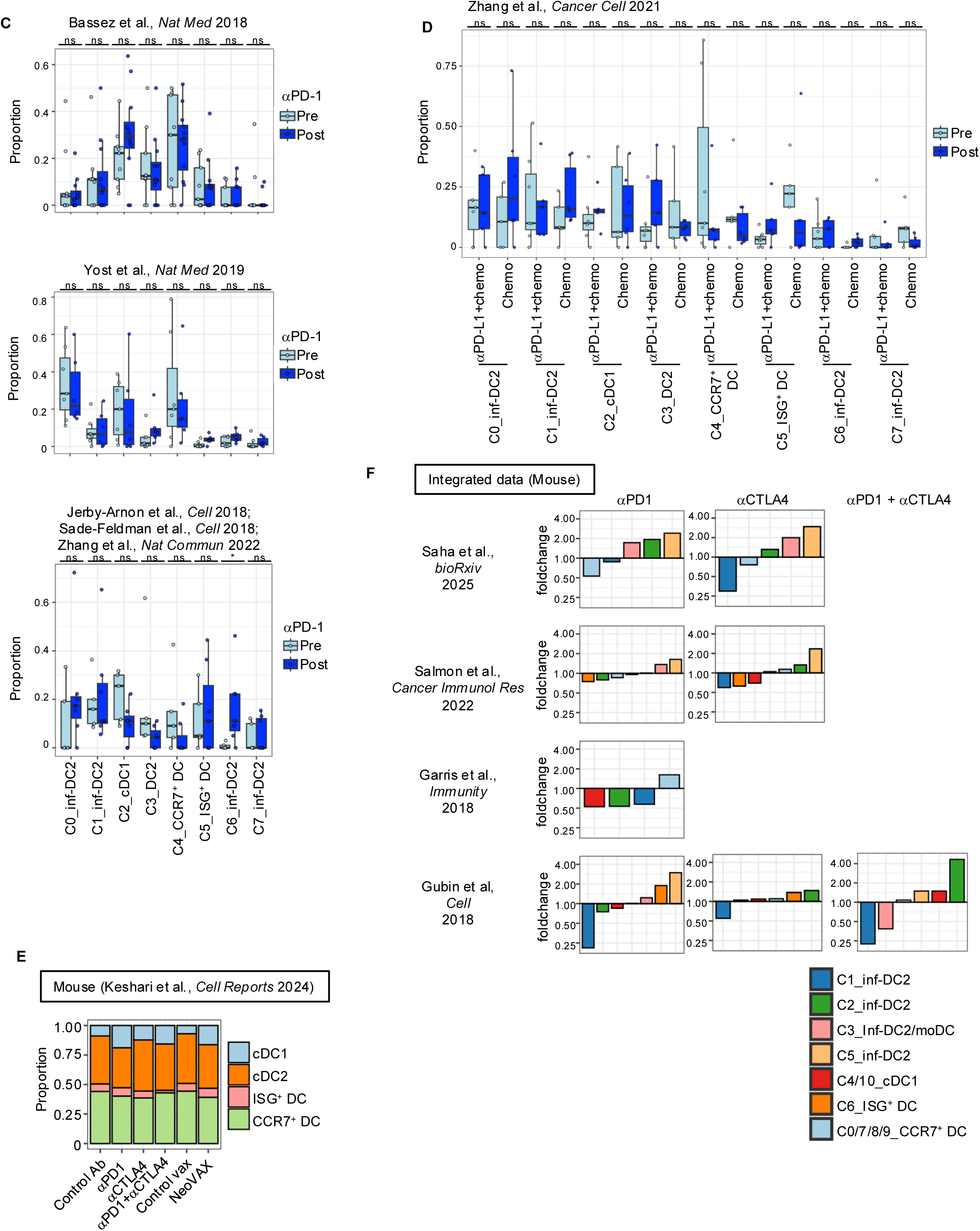
Transcriptional analysis of tumor-infiltrating DCs following immunotherapy across mouse and human datasets. **(A)** Dot plots summarizing the number of cells within each DC subtype across treatment conditions in the scRNA-seq datasets from Keshari et al.(*40*) (left), the integrated murine scRNA-seq dataset generated in this study (middle), and the integrated human scRNA-seq dataset generated in this study (right). Only datasets with available treatment annotations were analyzed. **(B)** Bar plots showing the number of significantly upregulated and downregulated DEGs across treatment conditions and DC subtypes in the scRNA-seq datasets from Keshari et al.(*40*) (left), the integrated murine scRNA-seq dataset generated in this study (middle), and the integrated human scRNA-seq dataset generated in this study (right). DEG analysis was performed by comparing each treatment condition with the corresponding control treatment groups. Only datasets with available treatment annotations were analyzed. **(C)** Box plots showing the proportions of DC subtypes comparing pre- and post-αPD-1 treated samples from breast cancer(*78*) (top), basal cell carcinoma(*79*) (middle), and melanoma(*80–82*) (bottom). The lines within the box and whiskers indicate the median and quartiles, respectively. P-values were calculated using one-way ANOVA. ns: not significant. **(D)** Box plot showing the DC composition in breast cancer patients treated with combination therapy (αPD-L1 plus chemotherapy) or chemotherapy alone(*83*). The lines within the box and whiskers indicate the median and quartiles, respectively. P-values were calculated using one-way ANOVA. ns: not significant. **(E)** Proportional composition of DC subtypes defined by reanalysis of scRNA-seq data from Keshari et al.(*40*) in murine tumors across treatment conditions. **(F)** Comparison of DC subtype fold changes across treatments and studies included in the integrated murine scRNA-seq dataset.

